# Ligand GA: a genetic algorithm for automated protein inhibitor design

**DOI:** 10.1101/2021.10.11.463970

**Authors:** Gordon Chalmers

## Abstract

Ligand GA is introduced in this work and approaches the problem of finding small molecules inhibiting protein functions by using the protein site to find close to optimal or optimal small molecule binders. Genetic algorithms (GA) are an effective means for approximating or solving computationally hard mathematics problems with large search spaces such as this one. The algorithm is designed to include constraints on the generated molecules from ADME restriction, localization in a binding site, specified hydrogen bond requirements, toxicity prevention from multiple proteins, sub-structure restrictions, and database inclusion. This algorithm and work is in the context of computational modeling, ligand design and docking to protein sites.

## Section 1: Introduction

The process of drug development is lengthy, tedious, and time consuming. Computer aided drug design (CADD) and structure based drug design (SBDD) are important methods in searching for viable small molecule drug candidates and protein function inhibitors. The advent and use of machine learning (ML) [1], AI techniques including neural networks [2], and large computing resources and storage are indispensable in modern day drug development. Genetic algorithms [3] and evolutionary algorithms [4] in general are a component of machine learning and are effective search algorithms.

The drawback of current day CADD or SBDD, however, is that limited databases are used in screening potential small molecule or fragments and in modifying their structures. Given a number of total heavy atoms and the atomic content, such as organic elements, the count of possible molecules can be exactly determined.^1^ This number can be considered large even with molecules of 100 atoms. The space of organic molecules made from {C,N,O,P,S,F} consisting of 100 atoms has a size of the order 10^80^ molecules. In contrast, public databases are drastically smaller. The E4C Exscalate4Cov project [5] uses a database of 10^13^ small molecules and fragments. PubChem [6] has 10^8^ molecules, Zinc [7] has 10^8^, DrugBank [8] has 10^6^, CSD [9] has 10^7^, ChemSpider [10] has 10^8^, ChEMBL [11] has 10^7^. These different databases not only have molecular structures, but in depth and different information about the molecules, drugs, their interactions, and availability which can be used in inhibitor design. If a desired small molecule in these databases is not close to any in these databases or in any of the databases, including proprietary ones, then the ideal candidate or a candidate will be missed in conventional drug design methods.

The ML approach presented in this work fundamentally does not use a database. In principle the GA will search the entire space of molecules for an ideal drug candidate. The primary limitation, as with all methods, is the finite amount of computing resources (e.g. CPU’s). GA’s are known for a good search if the mutation and crossover functions are appropriately chosen. Those in Ligand GA are modeled on physical chemical reactions as occurring in nature or in the lab. Information from databases however, can be included in the Ligand GA search for different purposes.

The software presented in this article fits into the initial stages of screening, hit and hit to lead identification, within the standard drug design methodology in 1. Furthermore, useful information that aids in the synthesis can be found in post processing the Ligand GA output: The GA iterations are stored as the GA executes, and from the different molecular modifications of mutation and crossover from one iteration to the next, possible synthesis pathways are generated. ***A link is given at the end of this paper; the download site has the software, documentation, and several examples. Much of the output from the runs in the examples was not included due to its size, but the necessary scripts for easily re-running are***.

The molecular representation is grounded in the use of the SMILES textual representations [12]. Molecules in the GA are textual chromosomes and Corina Classic^2^ [13] is used to convert these to geometric coordinates in mol2 or pdb files. SMILES has the feature that it contains the connectivity (adjacency matrix) of a graph and also the chirality of the chiral centers specific to valence 4 atoms and cis-trans of double bonds (i.e. stereochemistry)^3^. This is sufficient bonding information to create Corina energy minimized pdb or mol2 structures. These molecules may differ slightly in coordinates from an Amber [14] force field minimized structure. The flow of Ligand GA is described in Figure 16, in the section describing the design of the GA.

The docking software CCDC GOLD [15] is then used to find a set of docked structures (poses) of the small molecule in the region of the protein binding site. This is done for each stereoisomer. GOLD is a genetic algorithm with multiple scoring functions to use based on ligand protein interactions that takes into account the rotatable dihedral angles, flexible rings, protein side chain flexibility, water solvents, and more.^4^

The software is MATLAB [16] based. It calls internally the proprietary Corina Classic molecular geometric construction software and the CSD GOLD docking software as external components. It also uses MGLTools [17], and is setup to optionally use the AutoDock Vina [18] docking software. Ligand GA is designed to be easily used with many parameters defaulted but tunable to molecular goals.

The reliability of generation of pdb and mol2 files, particularly at chiral centers, has been studied in various papers, and Corina is comparatively one of a couple most reliable. Docking software have also been comparatively compared and CSD GOLD is efficient and very reliable, e.g. [19].

There has been use of genetic algorithms in molecular design [20,21]. These earlier GA’s use site specific fragment libraries based that reduce computational costs at the expense of a limitation on the search space, and use of a chemically restricted set of mutation and crossover functions. This GA is general with atomic modifications and pseudo-atom (e.g. fragments), and it has been generalized to multi-protein systems (Ligand Multi-Protein GA) since first submission with a multi-objective genetic algorithm (as discussed in the download documentation folder), although at higher computational cost. Also, this work uses AM+MN Corina Classic and CCDC GOLD docking, not OpenBabel and AutoDock which commonly fail in complicated pharmaceutical drug-like molecules. No comment can be made in comparison to any unpublished proprietary machine learning software.

Synthesis of the computationally generated molecules is important, but this broad and in-depth area is outside the scope of this work. Apart from visual inspection, inclusion of constraints in Ligand GA, and editing generating molecules, The MIT research program Machine Learning for Pharamaceutical Discovery and Synthesis Consortium (MLPDS) https://mlpds.mit.edu is dedicated to analyzing structures for synthesis planning, likelihood, and other related information. There is a site https://askcos.mit.edu that can also be used to a general run through of molecular results from Ligand GA.

This paper is organized as follows. Section 1 contains the introduction. Section 2 describes the principles of Ligand GA software design and 2 applications, COX-2 and Mpro of SARS-Cov-2 main protease (aka 3CL-protease), in the purpose of applying the software to specific inhibitors of interest.

Section 3 explains the overall GA software design. Section 4 concludes with a discussion of the software and its use.

## Section 2: Methodology of Ligand GA, Design, and Examples

Ligand GA is built with a specially designed GA with molecular modifications to forward an algorithmic evolution of a set of molecules to scan the solution space for optimality. This optimality is based on docking and binding of small molecule inhibitors to protein binding sites – active, secondary, or any site.

The protein-ligand interactions are used within CCDC GOLD to define and quantify a bound fitting of the ligand to the protein. A more accurate measure would use the energy of the protein-ligand complex in solvent –(minus) protein in solvent energy –(minus) ligand in solvent energy. This difference is a relative measure of the complex interaction taking into account the solvent and its displacement, not just the measure of protein-ligand interactions without surface solvent effects, which potentially could be relevant given the molecule surfaces and molecular occupancy in solvent.

The software is designed to include requirements, such as structural or chemical constraints, of the molecular goal. This section describes salient features and advantages of this approach to typical CADD or SBDD algorithmic methods and software. Two applications of Ligand GA usage are presented: generation of modified Aspirin and modified Simeprevir molecules. These 2 FDA approved molecules, each with 3 examples, were chosen to illustrate how Ligand GA can be used, what the output is, and in including of restrictions on the molecules.

The GA design is discussed in Section 3: Software. The methodology in the overall design of Ligand GA and its versality is demonstrated in the use of Aspirin or Simeprevir, with multiple results:

- Absorption, Distribution, Metabolism, Excretion (ADME) orally ingested restrictions on small molecules – Aspirin
- Examining direct output – Aspirin
- Tuning of molecular modification parameters -- Aspirin
- Editing unrealistic output molecules to make realistic -- Aspirin
- Molecules within the same binding site but localized differently -- Simeprevir, in stereoisomers and localization within binding site
- Excluding unwanted functional groups – Aspirin and Simeprevir

In Section 3: Software, the types of restrictions are discussed in implementation,

- Amino-acid hydrogen bonding requirements
- Unwanted non-target protein binding, as is done in virtual screening with large datasets, to avoid the toxicity problem
- Database inclusion in molecular evolution
- Constraints on molecular properties such as substructure or branch/functional group exclusion, and in using modification probabilities that influence features such as number of rotatable bonds, or atom content, and short scripts to exclude specific molecular features in the GA evolution (all in this set is general for any Ligand GA usage)

### Orally ingested small molecule ADME requirements, Aspirin

First order ADME restrictions can generally be imposed in drug design by enforcing Lipinski’s Rule of 5 [22]. There are an expanded and particular set also from the Ghose filtering [23], Veber’s rule [24], RO3 rule of 3 [25]. Lipinski’s Rule of 5 states:

The molecular mass is ≤ 500 Daltons

The number of hydrogen bond donors is ≤ 5.

The number of hydrogen bond acceptors is ≤ 10.

The octanal-water partition coefficient log K_ow_ is ≤ 5.

An extension has that the number of rotatable dihedral bonds is ≤ some number, e.g. 10 in Veber’s rule or 3 in RO3. The three mentioned expanded heuristics also change the limits of donors/acceptors, mass and log K_ow_, and have bounds on the number of atoms and polar surface area.

This is achieved in the GA by including a penalty on the molecules in the fitness function. ADME restrictions using Lipinski’s rule and others in drug candidate screening are examined periodically in the literature as the set of approved drugs in data sets increases [26]. Modifications of Aspirin are made using Ligand GA, and this molecule is used to illustrate how ADME restrictions are implemented in the software for an orally ingested drug. The x-ray complex PDB 4PH9 is used instead of PDB 5F19 due to the non-acetylation of the Serine 531 with Ibuprofen attached to COX-2.

Figure 2 shows the effect of using an ADME soft restriction on the evolution of modified Aspirin binder fitnesses. The difference is in the restriction of ADME generating lower fitnesses than the unrestricted. The mean and best fitnesses do increase (these are negative due to minimization) but less so with the restriction due to constraining the evolution. This is due to constraining evolution by Lipinski’s ADME rules and rotatable dihedral bonds less than 7. Note that the best fitness does not monotonically decrease; this is because the docking process is not absolutely deterministic and a GA docking result can give slightly different results even from the same starting molecule. These modified Aspirins have up to 2x PLP docking scores, satisfy ADME structural heuristics, and in general do not look like Aspirin. The reversibility or irreversibility of the inhibition was not investigated in the acetylation of Serine 531 as Aspirin does on COX-2 [27].

**Figure 1:**
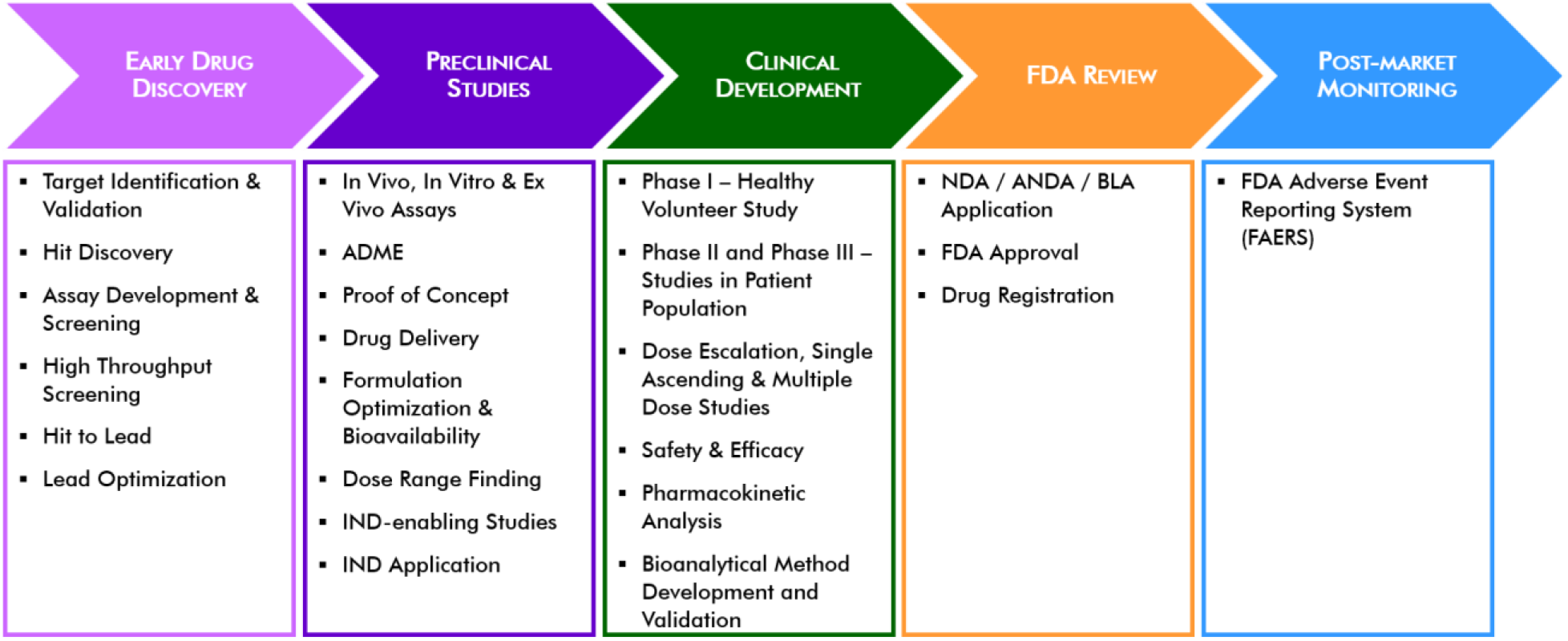
Drug Development flow. The Ligand GA software fits into the early drug discovery step and somewhat in synthesis.

**Figure 2:**
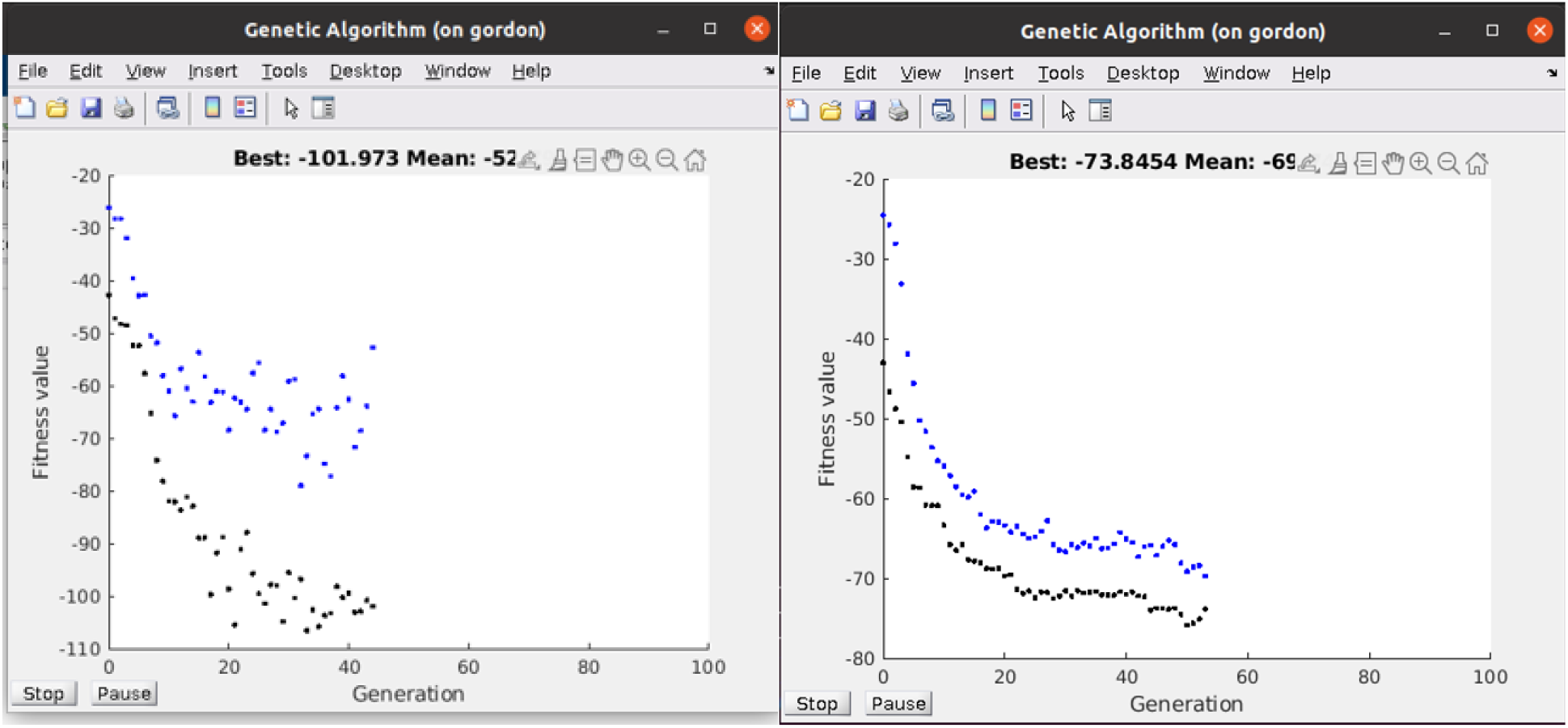
Fitness of population in molecular evolution of modified Aspirin. This pop-up during the program execution shows the state of the population of 50 molecules without ADME and with ADME restriction of a modified Aspirin run. The runs were initialized with 50 identical Aspirin molecules. Note that the restriction to Lipinski Rules of 5 and the total rotatable dihedral angles ≤ 7 are not binding as much, but still twice than Aspirin. PDB ID: 4PH9.

### Direct output of modified results, Aspirin

Aspirin (PubChem CID: 2244) has the isomeric form,

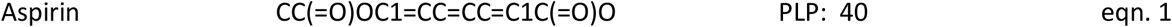

with GOLD PLP score of 42. It’s structure and docking is shown in Figure 3. The binding site is well in the protein A chain of the COX-2 dimer and contains SER A 531. A protein surface in the image obscures the image beyond visibility. The active residues of the protein in the GOLD binding of aspirin are: PHE206 PHE210 GLY228 VAL229 VAL345 TYR349 VAL350 ASN376 ILE378 PHE382 TYR386 SER531 GLY534 LEU535

**Figure 3:**
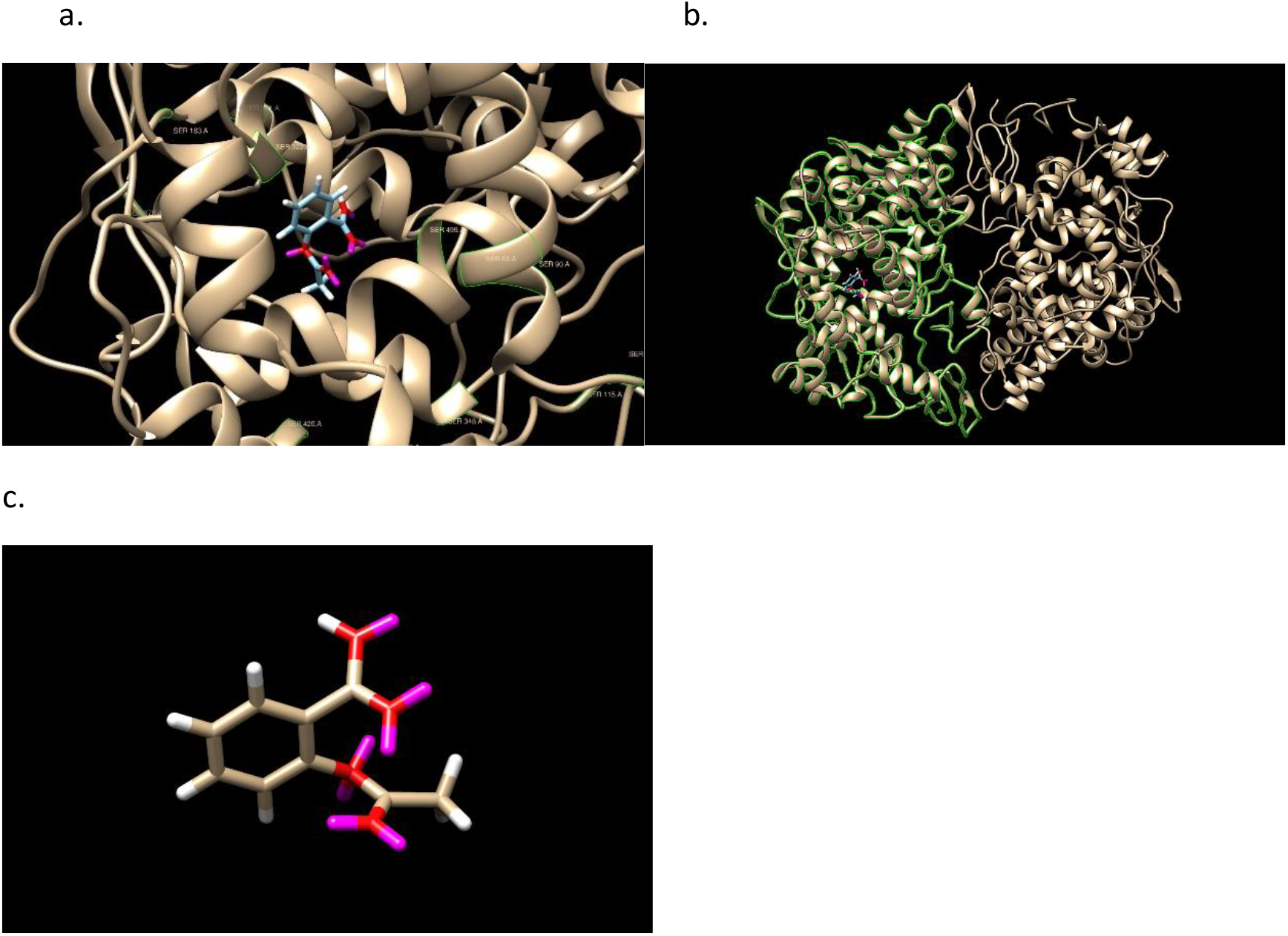
Aspirin docked into 4PH9 and Aspirin in the docking pose. The monomer A is shown in the second. GOLD PLP score 42. Note that SER531 in the pdb file is 499 in Chimera due to a numbering offset. Aspirin is next to SER531. SER is labeled in green. Please use a Zoom to see labels.

First, 3 -*conservatively-*chosen modified Aspirins directly from the Ligand GA output with a soft restriction^5^ of 5 rotatable dihedral angles, not 7, is given. Then in a different more specialized run with tuned mutation probabilities and atomic content, a high scoring example is given which is found after editing one of the output molecules.

The 3 modified Aspirins have improved PLP and from the direct output:

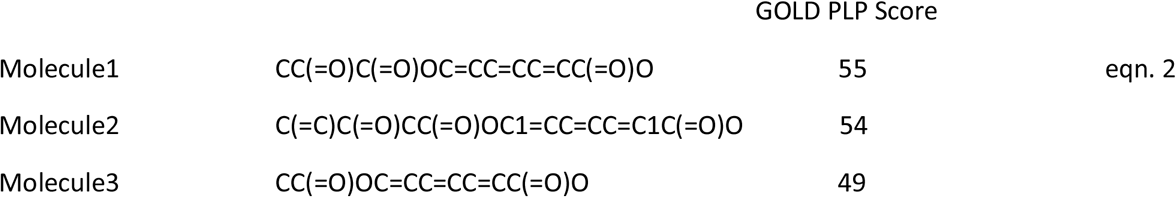

The oxygen number, 4 or 5, is about the same, 4, as Aspirin. The binding site of COX-2 that attracts Aspirin is known to be hydrophilic. The cis-/trans-stereochemistry of these molecules (1^st^ and 3^rd^ have 8 stereoisomers) is illustrated in Figure 4.

**Figure 4:**
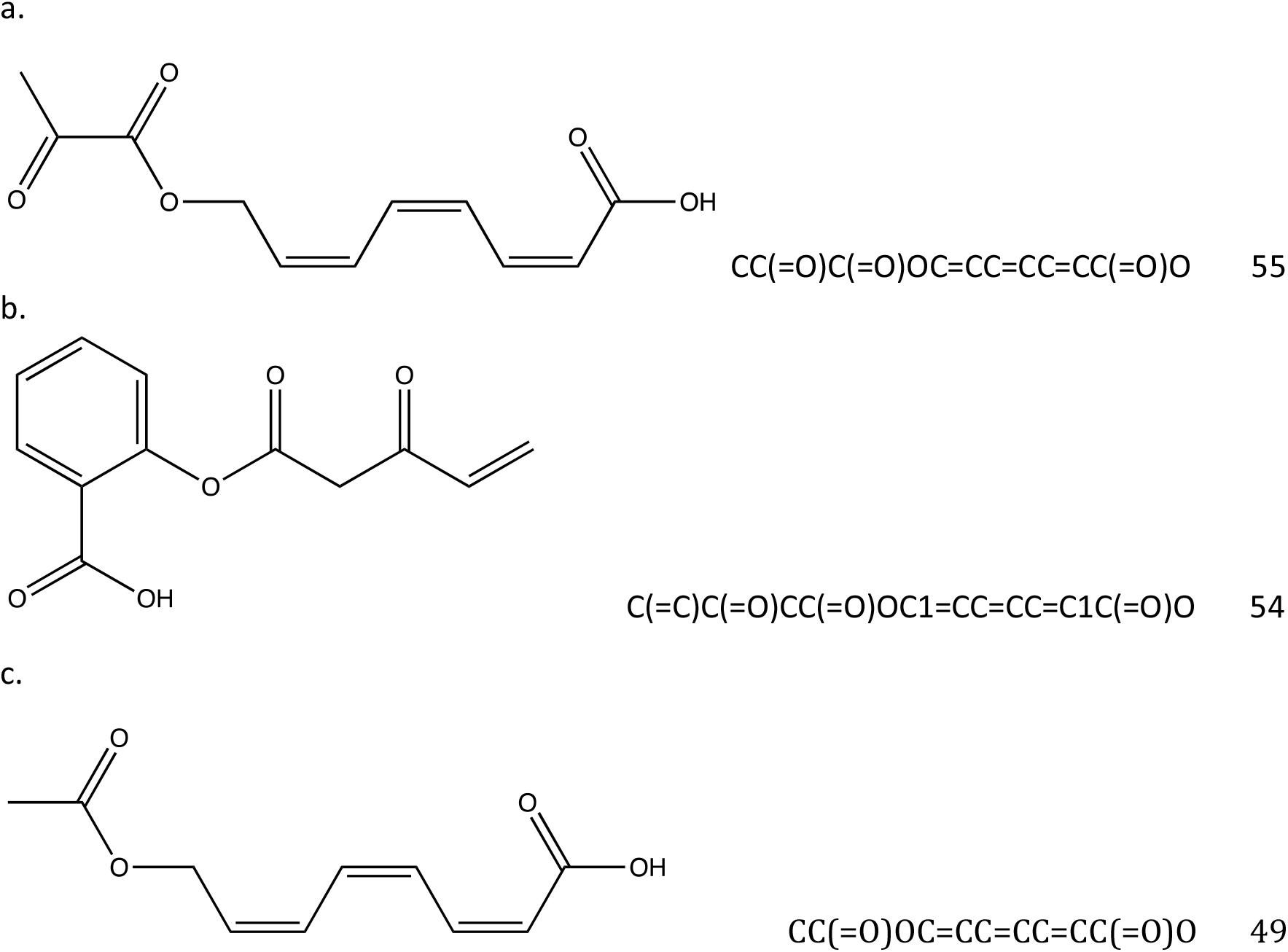
Structures of three modified Aspirins. They have 7,6,6 rotatable dihedral bonds and 16,17,14 heavy atoms.

For each of the 3 molecules, the highest docked pose scores are 55, 54, 49 and the docking is pictured in Figure 5. Figure 6 has an overlay of the different molecules in the protein binding site. From the docking output files, the active protein residues used by GOLD in its docking GA calculation are: PHE206 PHE210 GLY228 VAL229 VAL345 TYR349 VAL350 ASN376 ILE378 PHE382 TYR386 SER531 GLY534 LEU535

**Figure 5:**
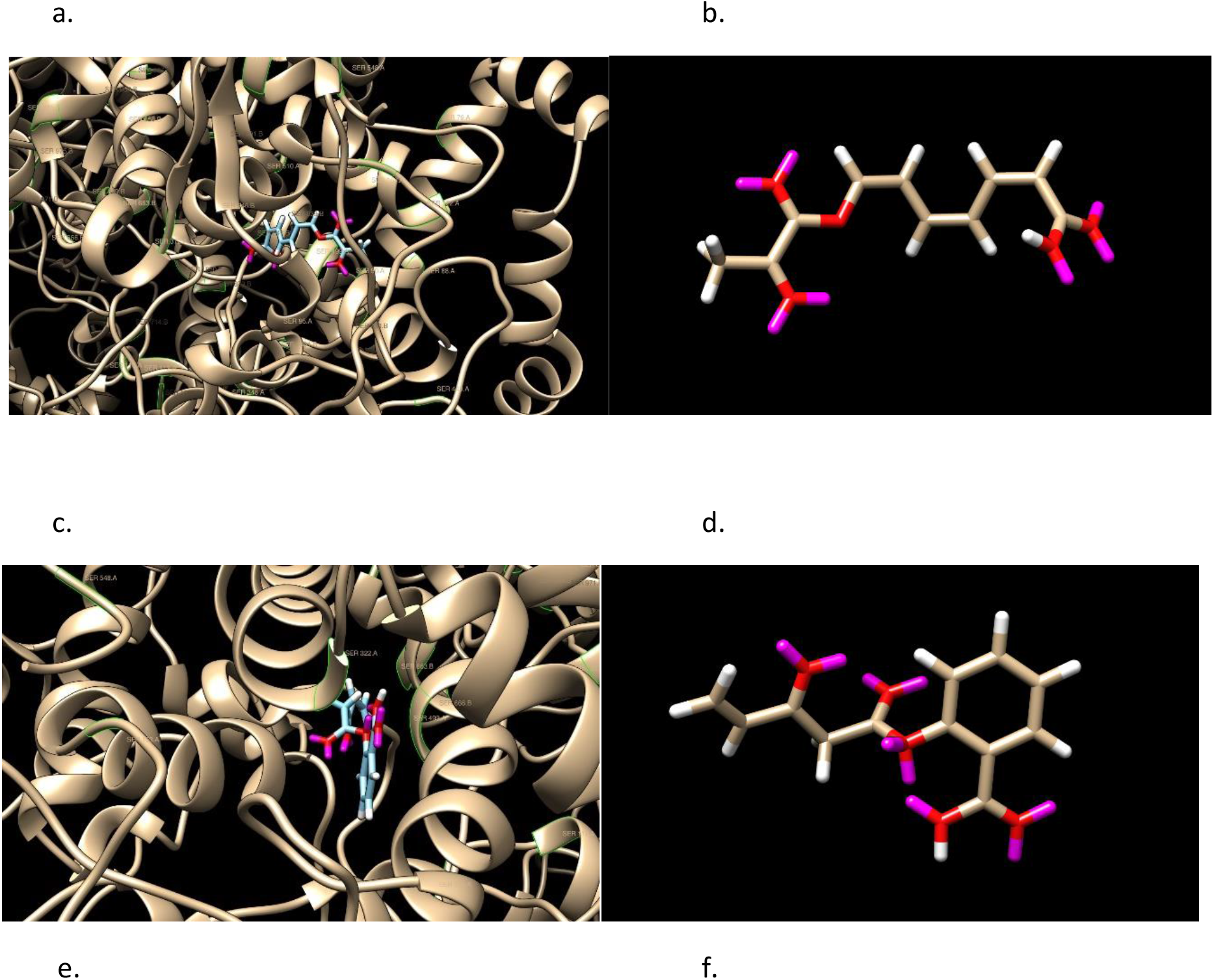
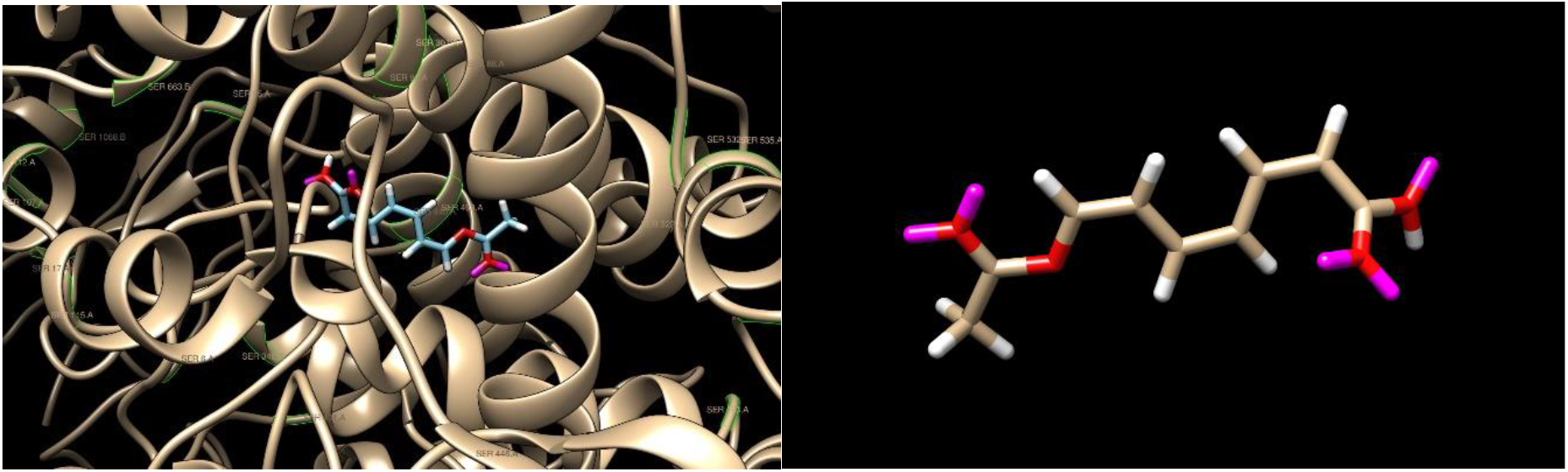
Docked modified Aspirin examples and images without the protein 6PH9 surface shown. GOLD PLP scores are 55, 54, and 49. Aspirin has a score of 42. SER is labeled in green.

**Figure 6:**
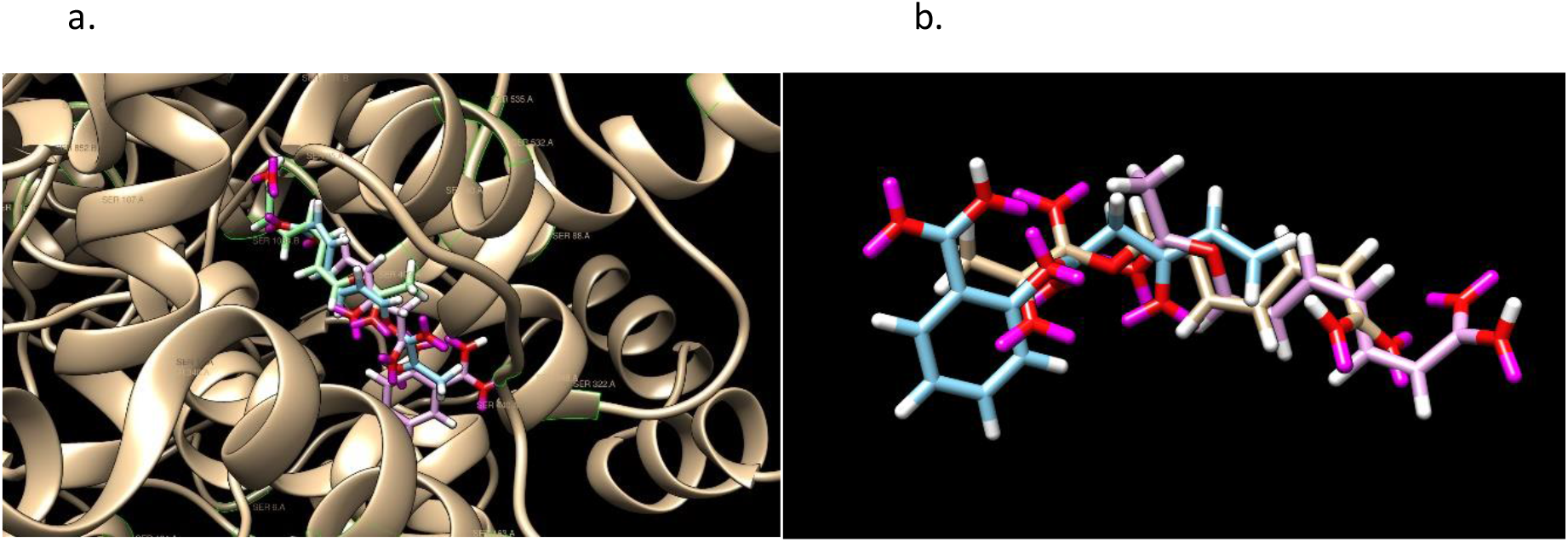
Superimposition of Aspirin and 3 modified Aspirins. Pink, green, tannish are in order of the listed molecules. Aspirin is bule. SER is labeled in green.

The set contains Serine 531, which acetylates when Aspirin binds and makes an irreversible inhibition. There are also multiple hydrogen bonds, not illustrated.

The 3 modified and Aspirin are shown superimposed in Figure 6.

The acetylation removal or requirement could be enforced of modified molecules, such as Aspirin, in the GA evolution by a geometric distance constraint from atom pairs between the small molecule and particular amino acids, such as Serine.

### Editing Ligand GA output molecules: Aspirin

The final example of Aspirin is shows how user interaction in post-processing can improve the result. In a hour run with 30 CPU’s, 634 molecules were generated from an initial population of 100 identical Aspirin molecules. SoftLipinski ADME restrictions were used in addition to C, N, O, F atoms. The mutation probabilities [3,1,2,1,1,3,1,1,.02,.02,.02,.02] were chosen to limit the increase of rotatable dihedral angles and also to minimize the use of single-triple and double-triple bond transitions; the decrease of these mutations also increased the performance of the calculation. One of output molecules has a PLP score of 96,

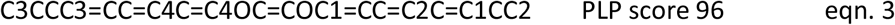

shown in Figure 7. This is a high score in comparison to Aspirin 42, but note that there are 2 enol ether groups make the molecule unstable in solvent. The molecule was edited in a variety of ways and using the fitness function on sets of possible changes, another molecule was obtained with a score of 91, shown in Figure 7,

**Figure 7:**
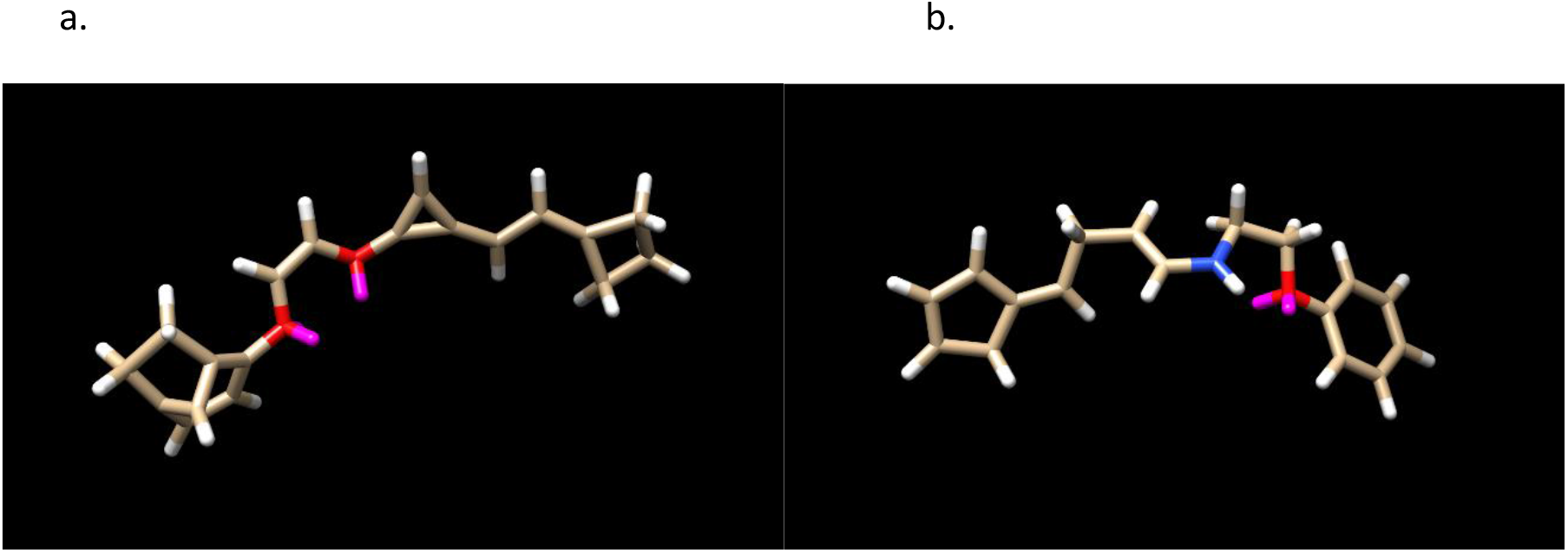
The LHS shows a high PLP scoring molecule in a tuned run with a population of 100. The RHS is an edit of the molecule having a PLP score of 90 and 2 stereoisomers. Both satisfy Lipinski’s Rule of 5.

**Figure 8:**
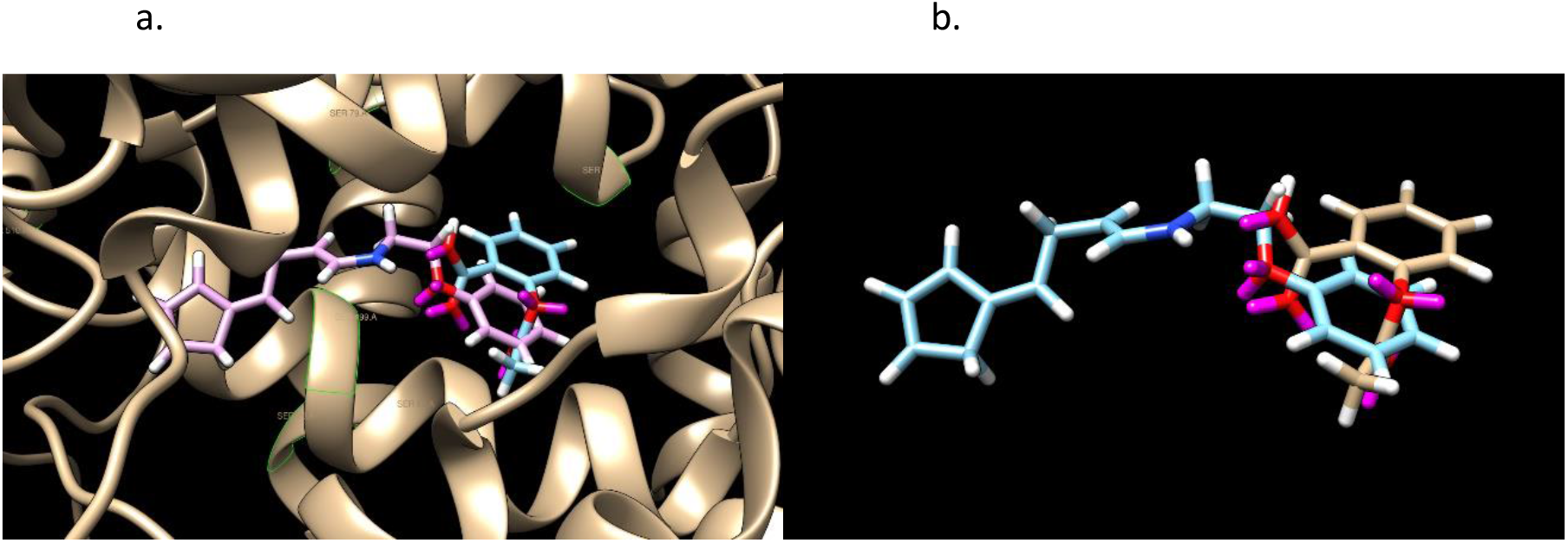
An overlay of Aspirin and the edited molecule in Figure 7. Note the orientation of the rings and the spatial overlap of the oxygens, 2 and 1 in the middle. There is one hydrogen bond from the generated molecule and two from Aspirin. GOLD PLP scores of 42 and 91.

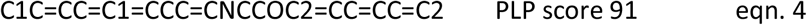

The procedure of editing the Ligand GA output demonstrates a realistic use of Ligand GA. After manipulating the molecule from Ligand GA, the 4-member, 3-member and the fused 2 rings were replaced with a 5-member and 6-member ring.

– In the starting molecule the 3-member homocyclic ring (unstable) has a double bond, suggesting to eliminate the 3^rd^ carbon. The ring did not provide additional docking stability and reducing the molecule size is relevant. Shortening the middle linear segment in most ways drastically reduced the docking score. As a result the double bond was kept, the adjacent O was replaced with an N, avoiding an enol ether, and shortening was not an option due to disproportionately large score decreases for an atom removal.
– Eliminating an atom between the 2 oxygens before changing one to a nitrogen would result in an unwanted -OCO-, which is also unstable. Thus, to eliminate the enol ether there, the double bond was changed to single and an -OCCO-was made, which is an ethylene glycol derivative.
– Next, the fused 2-rings eliminated 4 atoms and was replaced by an aromatic carbon ring. The 4-ring was changed, for a better score, into a 5-ring with 2 double bonds.

The molecule has 7 rotatable dihedral bonds; there would be 6 if the double bond from the 3-member ring was changed to a synthetically difficult triple bond but the score decreases to 82. There are no enol ethers and the molecule looks ‘normal’, being from joined aromatic 6-ring—ethelyne glycol derivative— 2 double bond 5-ring. These changes were selected from a combined set of various changes with the docking score and protein-ligand interaction in mind. The docking of the molecule appears prefer length in one direction, and the N and the O. In checking the rings, there doesn’t appear to be a difference (in the CSD GOLD calculated result) if the 6-ring is a 5-or a 6-having double or single bonds, with the aromatic 6-ring slightly higher on average in multiple identical docking calculations. The 5-ring on the other side prefers 2 double bonds in it. The N and O each are bonded to 2 carbons and changing the oxygens to nitrogens did not make a large difference in docking score but in possible stability of the molecule.

Aspirin has 13 heavy atoms and the designed one has 19; this is a 46% increase. Docking score is expected to increase with more heavy atoms, not necessarily linearly, but so is flexibility, which can be a drawback for a small molecule; the docking score increased 114% and the flexibility from 3 to 7. Aspirin and the designed one has 2 and 1 hydrogen bonds in these poses. While Aspirin has no chirality, this designed molecule has 2. Some drugs have large numbers of both rotatable dihedral bonds and molecular mass, and cannot satisfy small molecule ADME guidelines.

A feature of the Ligand_GA_Fitness_Function in this step is that it takes an input of a list of molecules expressed in SMILES and uses the GOLD configuration file for the docking. As a standalone function, and without any relevance to a genetic algorithm, it will create all of the stereoisomers for each molecule from the input (also if the SMILES expression is partially and isomeric and non-isomeric), creates the output directories in the ligand_dir directory, and docks all of them. It is a useful function for the iterative process of taking an output molecule from Ligand GA and making it more realistic.

In these runs, there are generally molecules with >50% more higher docking score,

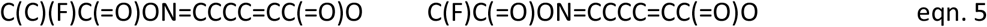

which satisfy ADME requirements and number of dihedral angles less than 7 or 8, but which contain no rings. Without ADME restrictions, molecules with scores up to 3x that of Aspirin were found and many were of comparable mass to Aspirin (scaling does not match 3x).

Ligand GA is designed to take in user criteria and produce computationally small molecules. The guiding requirement is high quantitative docking scores. The output is a set of unique small molecules.

### Stereoisomers, Structural localization in the binding site: Simeprevir

A very important point, not often addressed in docking studies, is that the binding site of a protein is not just a region of the surface of the protein. The site will have a geometric terrain of hills, holes, valleys, and ridges. This can be found in a visual inspection of the site in Chimera [28], VMD [29], or PyMOL [30], and particularly in Nanome [31] with an Oculus Quest 2. There are many different high binding molecules that will bind in this region, but which have different physical localization and chemical binding into the terrain of the site. The interaction of a protein’s surface as a substrate is in the specific local details of the terrain and its amino acid content within the substrate and the protein binding site. Simeprevir (PubChem CID: 24873435), with trade name Olysio, is an FDA approved therapeutic for treating Hepatitus C [32]. It gained interest as a repurposed inhibitor of the main protease Mpro (3CL-Pro) of SARS-Cov-2 early on in the 2020-21 pandemic at the binding site of Boceprevir [33]. The molecule has mass 750 Daltons, hydrogen donor count of 2, acceptor count of 10, log K_ow_ of 5.16, and 8 rotatable dihedral angles. It violates Lipinski’s Rule of 5 and dihedral angle count in 3 of the conditions, and borderline 4. Simeprevir is also different in that it has a large ring with many bonds.

3CL Mpro of SARS-Cov-2 has a quantitative structural similarity to the class of NS3/4A proteases, in particular to that of the Hepatitus C virus (HCV) [34]. In the latter, the surface structure has features such as a shallow groove which makes it difficult to find small molecule inhibitors, and this part of the site has protein-substrate interactions relevant to HCV viral replication. The desired inhibitor clearly requires specification of amino acids in the target site for effectiveness, and Ligand GA can provide this, as is the case in irreversible inhibitor design that requires specific target amino acids.

A set of inhibitors of the SARS-Cov-2 Mpro has been investigated computationally, in docking, and in experiments to find synergistic combinations of known drugs to block Covid-19 [35]. Simeprevir ranks high in the list of molecules in AutoDock scores.

Figures 9 and 10 shows the stereochemistry and docking of Simeprevir to Mpro (3CL-Pro) of SARS-COV-2 and of a stereoisomer which has a 25% greater GOLD PLP docking score. The GOLD scores are 71 and 89.

**Figure 9:**
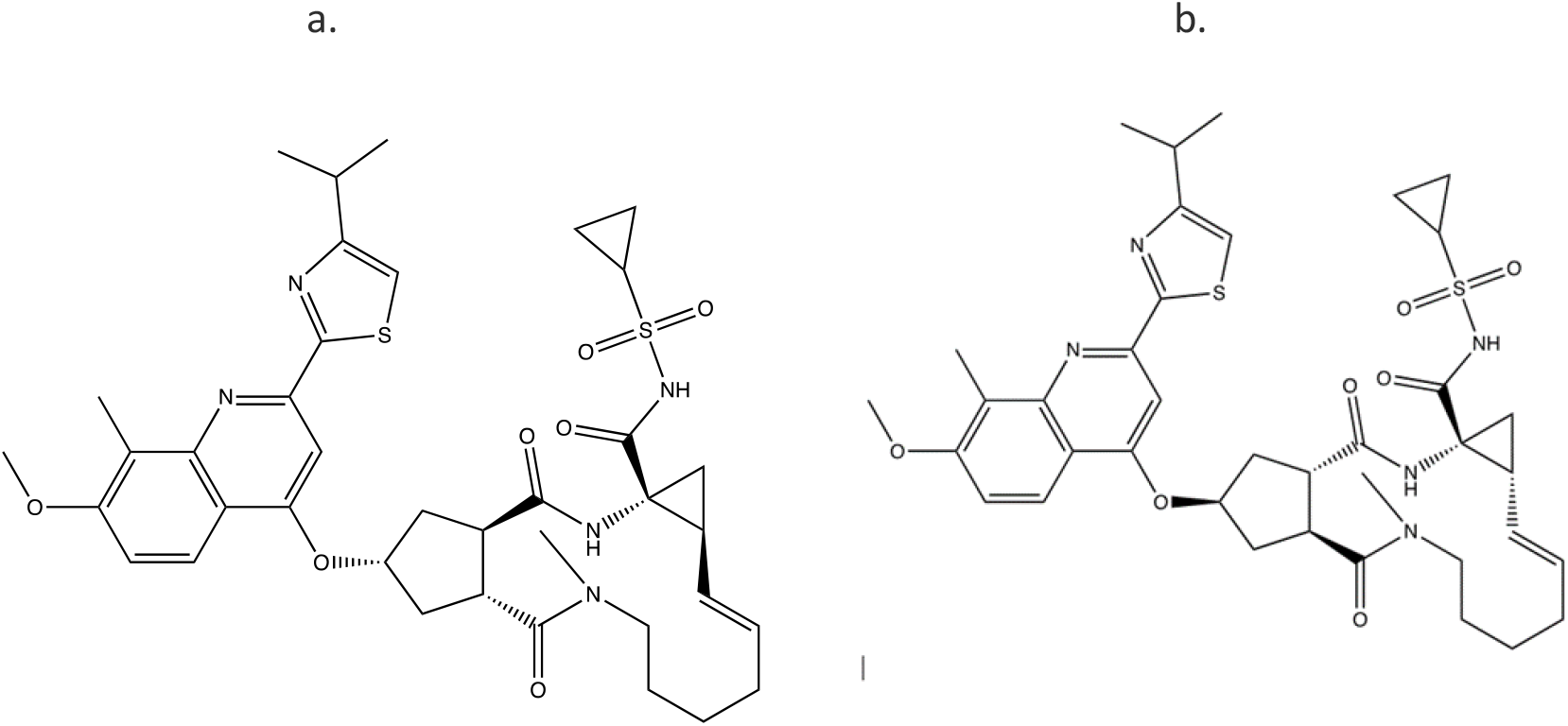
(a) Simeprevir, an FDA approved drug for treating Hepatitus C. It has been considered as an inhibitor of the Serine type of protease Mpro of SARS-Cov-2. GOLD PLP score is 71. The docked result used an isomeric chiral SMILES string. (b) Stereoisomer 5 of Simeprevir from Corina (5 label comes from the basis set ±, ±, ±, ±, ± of chirality of the stereocenters from a non-isomeric SMILES string). When bound to Mpro of SARS-Cov-2 its docking score is 48% higher than Simeprevir. GOLD PLP score of 89. Note that 4 out 5 chiral centers in (b) changed direction and structurally this molecule is different from the one in (a).

**Figure 10:**
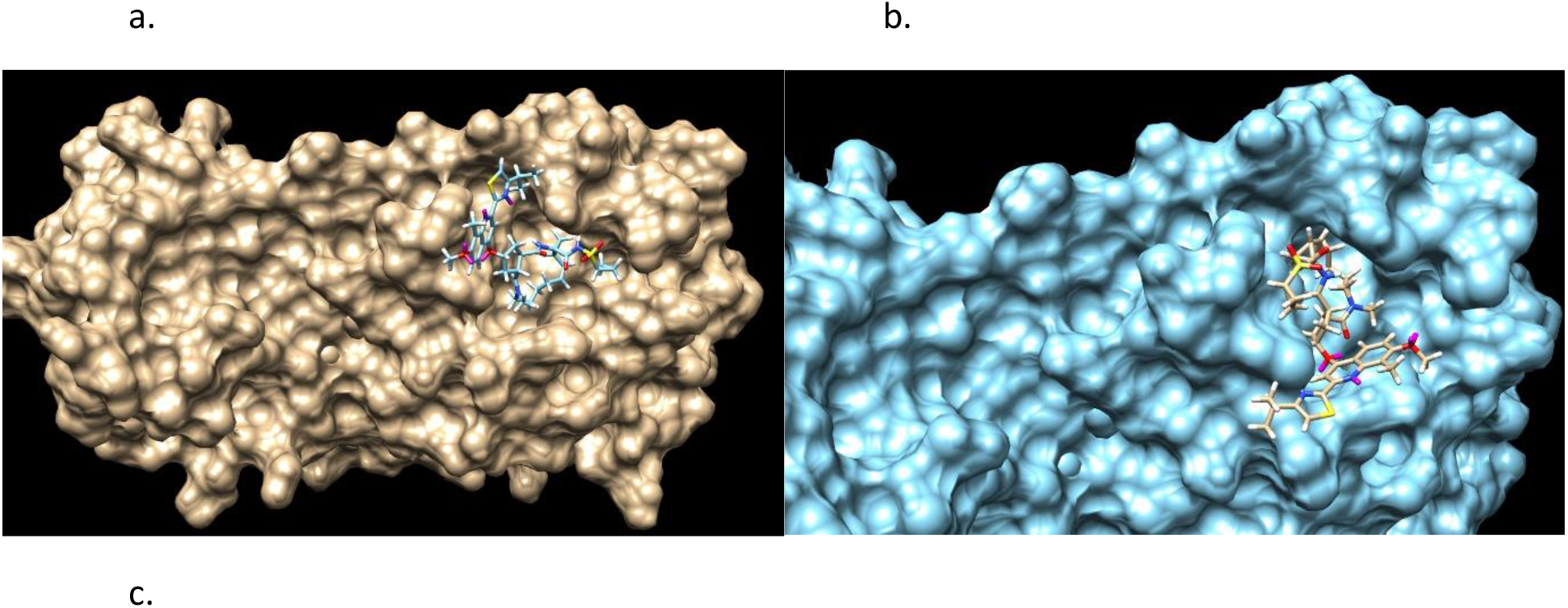
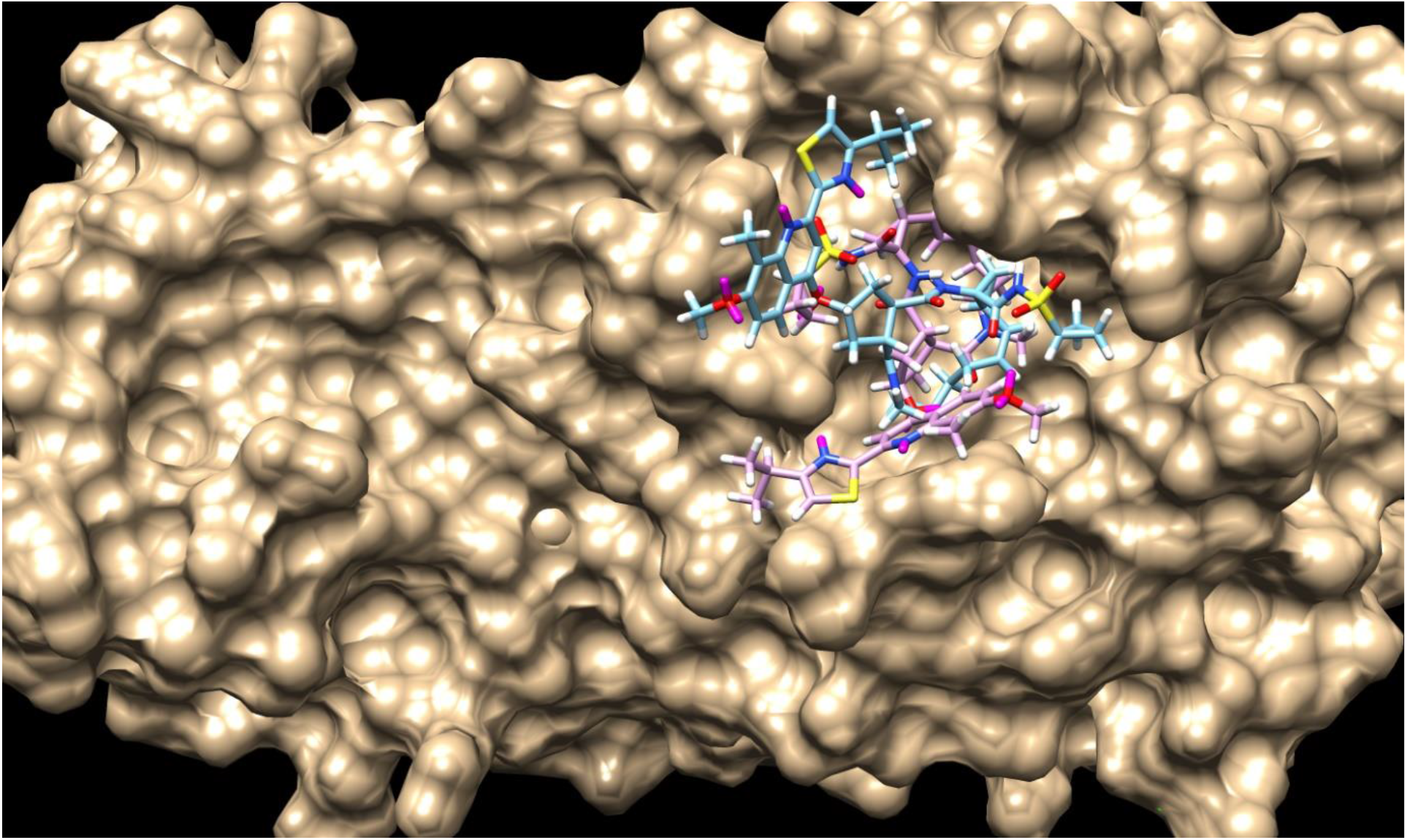
Simeprevir and Stereosimoer 5 of 32 at the BOC site in the SARS-Cov-2 Mpro main protease. GOLD PLP Scores are 71 and 89. The overlay shows the differences in localization. PDB ID: 6WNP. #5 is the pink ligand in the overlay.

*Importantly, Simeprevir was chosen in this presentation for the reason of it being a well-used drug and also because of the interest it has had as an inhibitor of the SARS-Cov-2 Mpro since the early period of the pandemic. There are an enormous number of inhibitor studies of the different proteins of SARS-Cov-2 [34]. The application of the Ligand GA software is general and this is only an example in a current global research area*.

Both bind to an active site which Boceprevir (PubChem CID10324367), a Calpain type of inhibitor, binds to in the SARS-Cov-2 Mpro main protease. Simeprevir has the non-isomeric SMILES form of

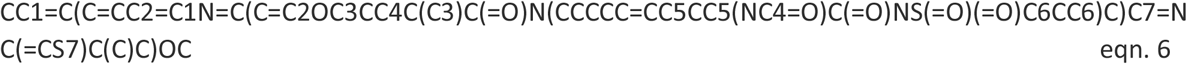

and the isomeric form,

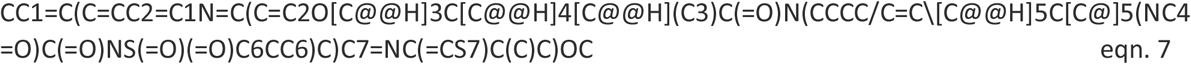

and does violate most of Lipinski’s rule of 5. The Simeprevir molecule isn’t ideal to illustrate the use of the software in modifying due to violation of ADME restrictions and flexibility, but it is of interest for multiple reasons.

### General, Simeprevir

Simepriver is a designed molecule to inhibit the function of HCV NS3A/4. It is already on the other side of Lipinski’s Rule of 5, yet it is an effective component of the FDA approved treatment of Hepatitus C. Due to design rigidity, generic modifications and repurposing of the molecule in order to treat other conditions are expected to break its stability or viability as a drug candidate. The following 3 examples out of hundreds of modified Simepriver show the aspects of modification using Ligand GA and the importance of applying simple constraints. These constraints are simply stated and also implemented essentially as short Matlab scripts called by Ligand_GA_Fitness_Function_GOLD_ADME.m.

Figures 11 and 12 are of a modified Simeprevir molecule bound to this site, of approximately 29% increased docking score. Two perspectives are shown. Due to Simeprevir violating the Lipinski’s Rules of ADME restriction, the number of stereoisomers increasing from 32 to 512 (default maximum in analysis), and computational resources, the ADME constraints with the initial population of 50 identical Simeprevir molecules was not included in the generation of the modified higher scoring molecules. The one in Figures 11 and 12 (docked) is the highest scoring in this limited run, but there are others. If the large ring is opened anywhere then the flexibility is drastically increased generally and thus the molecule is more flexible, which can be a problem. This molecule was chosen to present in the paper due to the interest in inhibitors of SARS-Cov-2 and it being a well-used drug. Molecular mutation probabilities in Ligand GA can be emphasized towards not opening this large ring but in shortening it by atom deletion or substituting, specific to this ring, which is relevant to this class of molecules. There were 349 and 210 generated in two separate 12 hour runs with 30 CPUs. The molecule is

**Figure 11:**
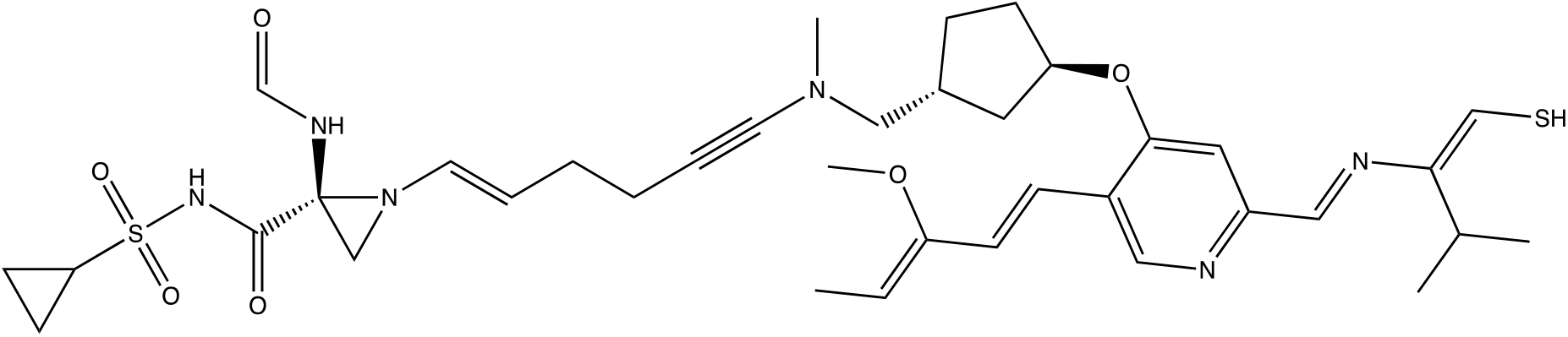
modified Simeprever example 1. An overnight run of Ligand GA generated 349 different canonical SMILES molecules, each of which have multiple stereoisomers. GOLD PLP score of 109. Most other molecules in this set also appear flexible due to the chances of opening the large ring in the GA molecular evolution.

**Figure 12:**
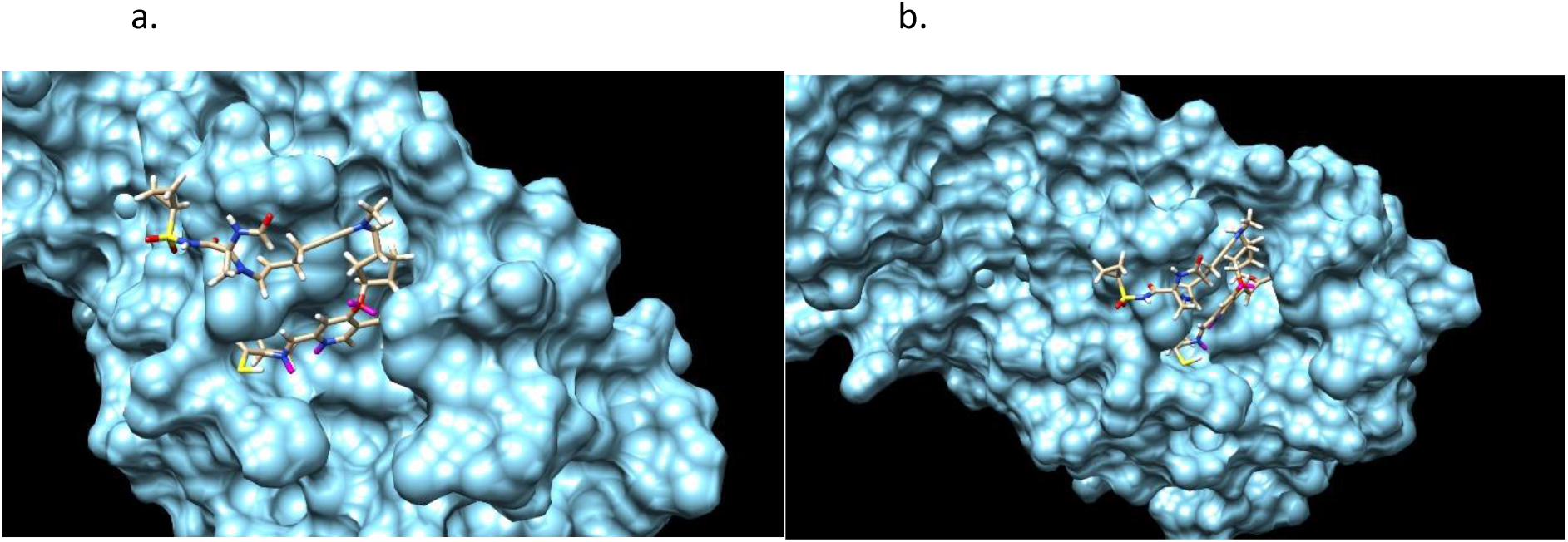
Modified Simeprevir stereoisomer 164 of 512. GOLD PLP Score 109. PDB ID: 6WNP. There are 349 different non-isomeric molecules from this run, each with multiple stereoisomers. 2 perspectives.

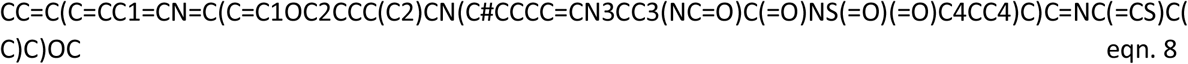

and has 4 rings, not 7 as in Simeprevir, and it also has 19 rotatable dihedral bonds. If a restriction is imposed that prohibits the opening of the large ring at the internal bonds then the flexibility won’t increase much in the generated set. This example also shows the relevance of a general total dihedral restriction, which was not used in this case, but could be included directly in almost zeroing out the mutation probability for OPEN_RING.

Another example of modified Simeprevir, and comparable GOLD PLP docking score 98, is

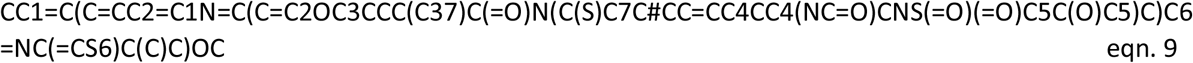

Figures 13 and 14 are of the 2^nd^ example molecule and docking. This molecule has fewer rotatable bonds than the previous, 11 in total and Simeprevir has 8. There are many structures generated in the output of the run and these are unconstrained examples. The output, as in design, could be modified for better structures. For example, is the carbon between the 2 nitrogens necessary for good binding? If not, then the flexible bond number is reduced by 2 and so is the mass. In the case of Simeprevir, which violates ADME requirements, Ligand GA provides starting points from the molecule output as does a large database in repurposing known molecules. These calculations are improved by adding more computing resources and additional constraints.

**Figure 13:**
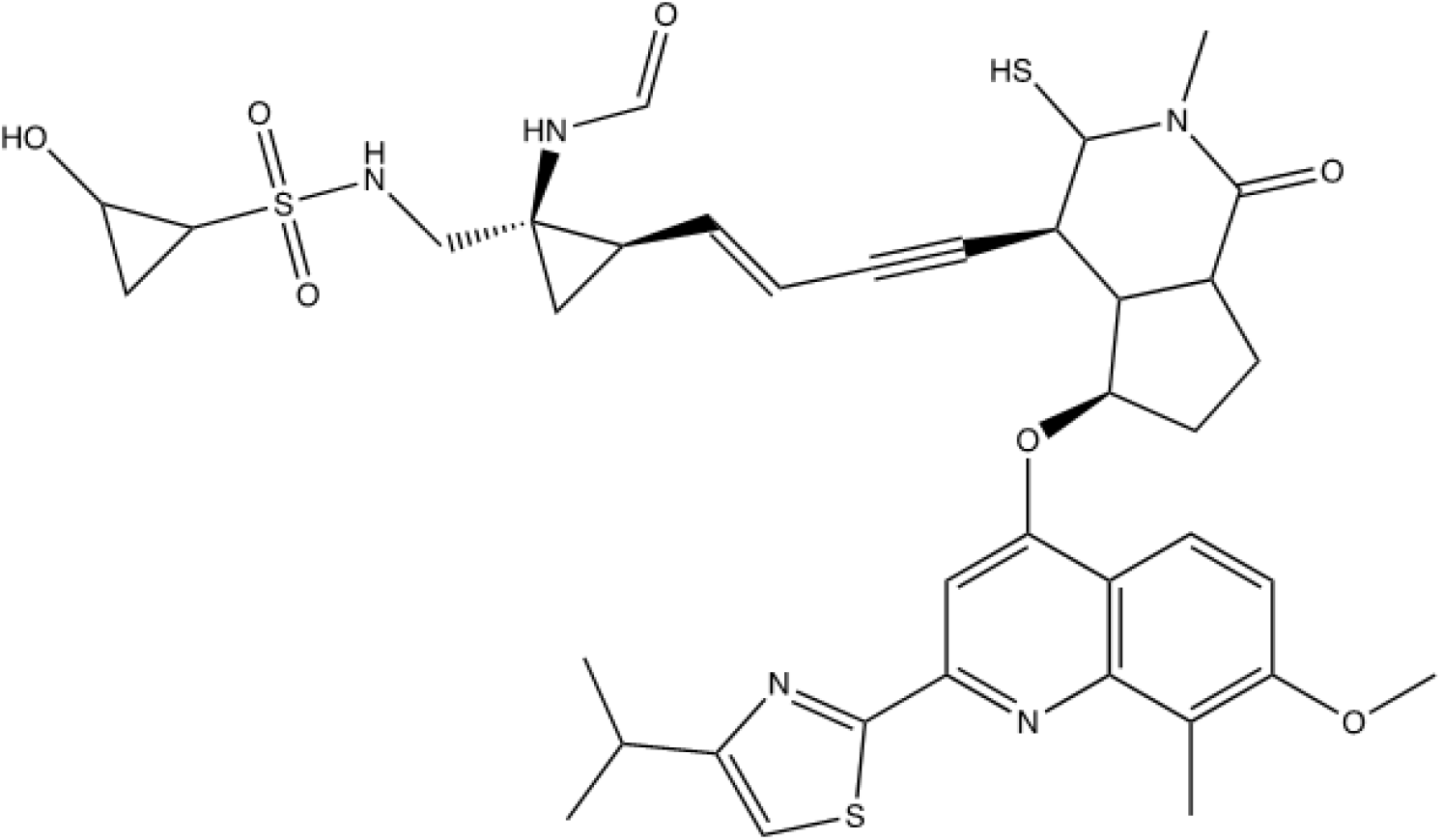
2^nd^ example of modified Simeprevir. GOLD PLP score of 98.

**Figure 14:**
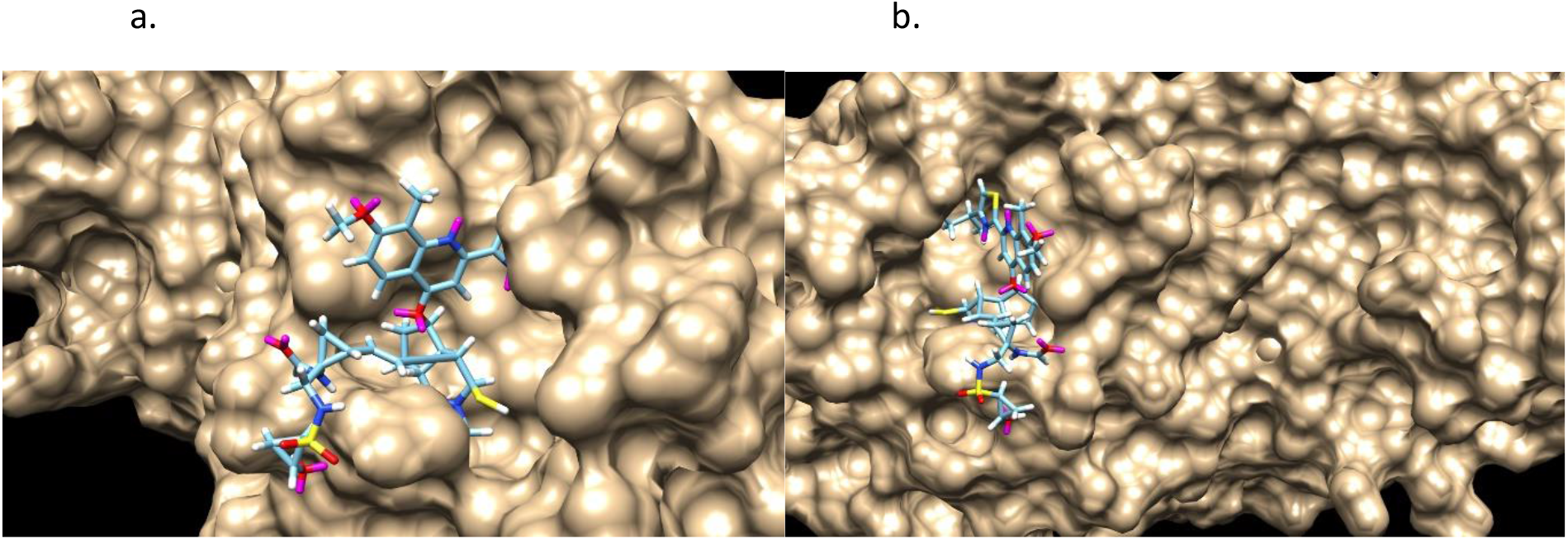
Modified Simeprevir stereoisomer 288 of 512. GOLD PLP score is 98. PDB ID: 6WNP. In this run there are 210 different non-isomeric molecules. 2 perspectives

The 3^rd^ example of modified Simeprevir is a much smaller molecule,

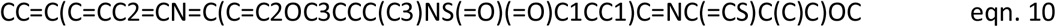

and this one satisfies more of Lipinski’s Rule of 5. It should be noted that the molecule in Figure 15 has an enol ether and thioenolate functional group, which would convert to a ketone and a thioketone in water and also an unwanted -SH. This compound is not stable in water, at one and maybe 2 end-points. The molecule has a GOLD PLP docking score of 77 and is illustrated in Figure 15, however it is unrealistic. Docking pictures, although fitting in the BOC site, are not included. These derivatives are minor and can be changed by hand if no automated replacement is done.

**Figure 15:**
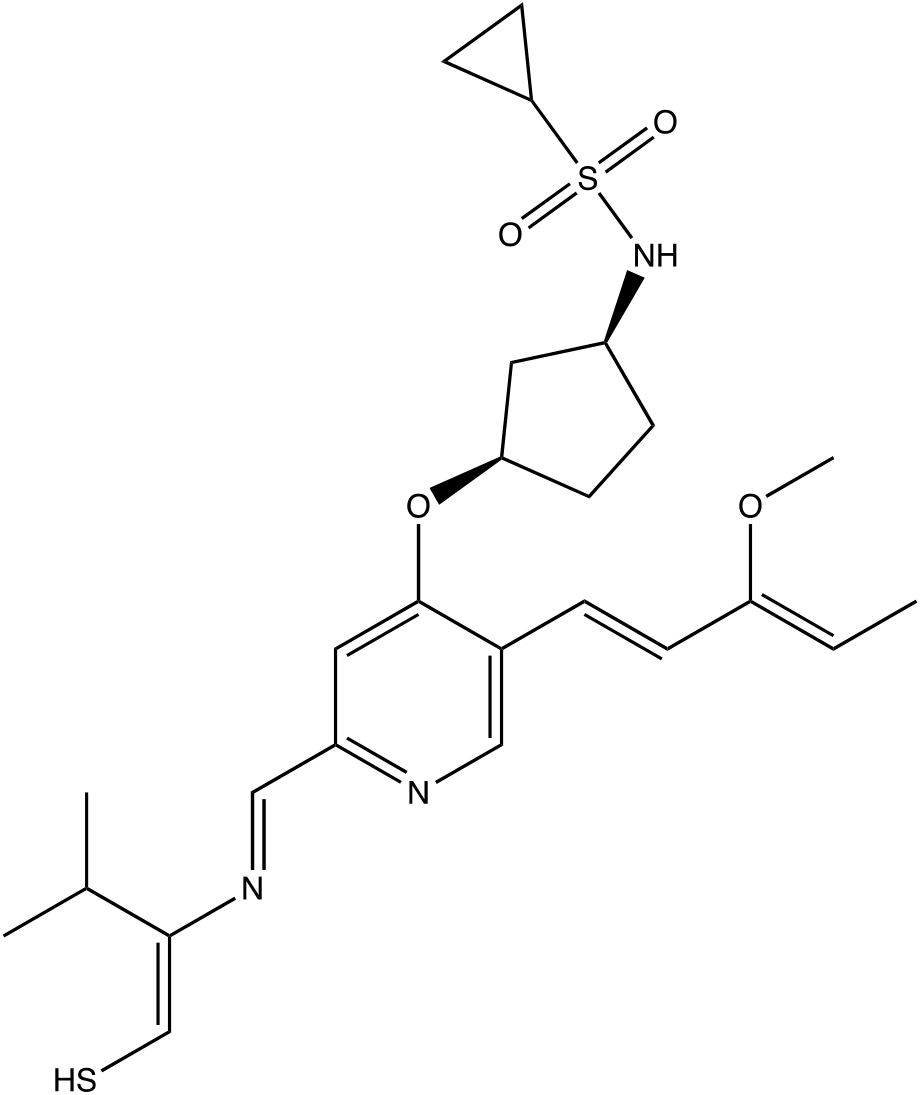
Example 3 of modified Simeprevir. It has 34 heavy atoms compared to 52 of Simeprevir. GOLD PLP score of 77.

This 3^rd^ example also illustrates why Ligand GA was designed to include constraints in the fitness function. An additional penalty can be included to penalize GA chromosomes, i.e. molecules, of having the -C(OC)=CC or -CCS at the end of branches. These penalties are from external scripts called from Ligand_GA_Fitness_Function_ADME.m; an example script is to penalize the molecule if a branch has either of these functional groups at end-points.

(The OPEN_RING function opens at SMILES ring points, not general ring bonds. The equivalence of different representations of the same molecule requires the OPEN_BOND function, which does this at bonds in rings. 7 rings means that if OPEN_RING is called out of all the molecular mutations then there is a 1/7 chance of opening the large ring. There are 38 ring bonds that could open, 13 of which open the large ring. OPEN_RING could preferentially open the large ring if the molecule mutation probabilities are uniform, and OPEN_BOND is not included in this version.)

In these examples, these molecules are docked into the site but cover different local regions. It is clear that these molecules are computationally designed because of the filling of the pockets within the site or in the covering of a valley or ridge. The specific localization of an inhibitor to a site can be required in the molecular construction if the specific mechanism of the binding site is known. This is indeed important in inhibitor construction because the very specific information of which amino acids should be inhibited chemically is included and not only the binding region.

To remind, as stated earlier, the HCV NS3/4A surface structure has features such as a shallow groove which makes it difficult to find small molecule inhibitors, and this part of the site has protein-substrate interactions relevant to HCV viral replication. An example source of large amounts of information pertaining to the SARS-Cov-2 proteins is at [5,37]. The creation of irreversible inhibitors such as Aspirin can also be enforced by specifying in the GA evolution a constraint of amino-ligand interaction, as in SER 531-Aspirin and its acetylation.

In the case of Simeprevir, the output contains hundreds molecules which fit differently in the binding region, even at higher but not much higher docking score.

### Hydrogen bindings

Ligand GA is designed to take in detailed user criteria. In addition to imposing ADME requirements or localized geometric information, the user can require the presence of hydrogen bonds to specific amino acids. Well placed H-bonds are important in inhibitory action and binding, as well as in the permeability of the molecule though different media and through different barriers such as membranes or cell walls. One well-placed H-bond can increase binding affinity by approximately 1.5 kCal or permeability up to 10x.

### Toxicity prevention

Lastly in this section is the question of toxicity or interference of other protein processes can be included. Virtual screening in commonly used CADD also include this in the amount of screening with multiple protein binding sites. If the target protein’s function from this site is to be blocked, and 5 other proteins and sites are not to be interfered by the small molecule, then a penalty arising from the unwanted binding can be included in the fitness function. GA will try to optimize a binder to one protein but then penalize the drug candidate if binding to the other protein sites. The computational complexity increases by a factor from the number of binding sites included in the fitness evaluation of the evolved molecule, and the software essentially does not change. However, the computational resources multiply. 30 processors now becomes 150 and so on.

All of these aspects have been included in the development of the Ligand GA software.

## Section 3: Software

In this section the Ligand GA software design is explained. The package download has documentation files, detailed presentations, and the commented code. There are also directories with the Ligand GA input and output files pertaining to its use in modified COX-2 and SARS-Cov-2 Mpro structures. Although database input was not used in these, the initial populations for the GA were identical copies of either Aspirin, Celecoxib, or Simeprevir. In general, any initial population can be used on any binding site, but initialization of the population is important for an effective search, and in any GA application, not just in Ligand GA.

The software has many ingredients. To simplify the construction and the use, the remaining section is broken into different important topics: *Ligand GA flow, Tuning, External software, Fitness function, Constraints, Databases, Crossover and mutation, Selection, Storage of results, Output and Post-processing*,

### Ligand GA flow

The logical flow of the Ligand GA is illustrated in Figure 16. With the 4 file/filename inputs and the parameters in the Lignd GA Global file set, the head function starts the process by evaluating the fitness of the individuals in the population. The steps in a GA are: initialization, evaluate fitnesses of the population, selection of the individuals, crossover and mutation. There is also the step of saving the information of all the individuals at each iteration.

**Figure 16:**
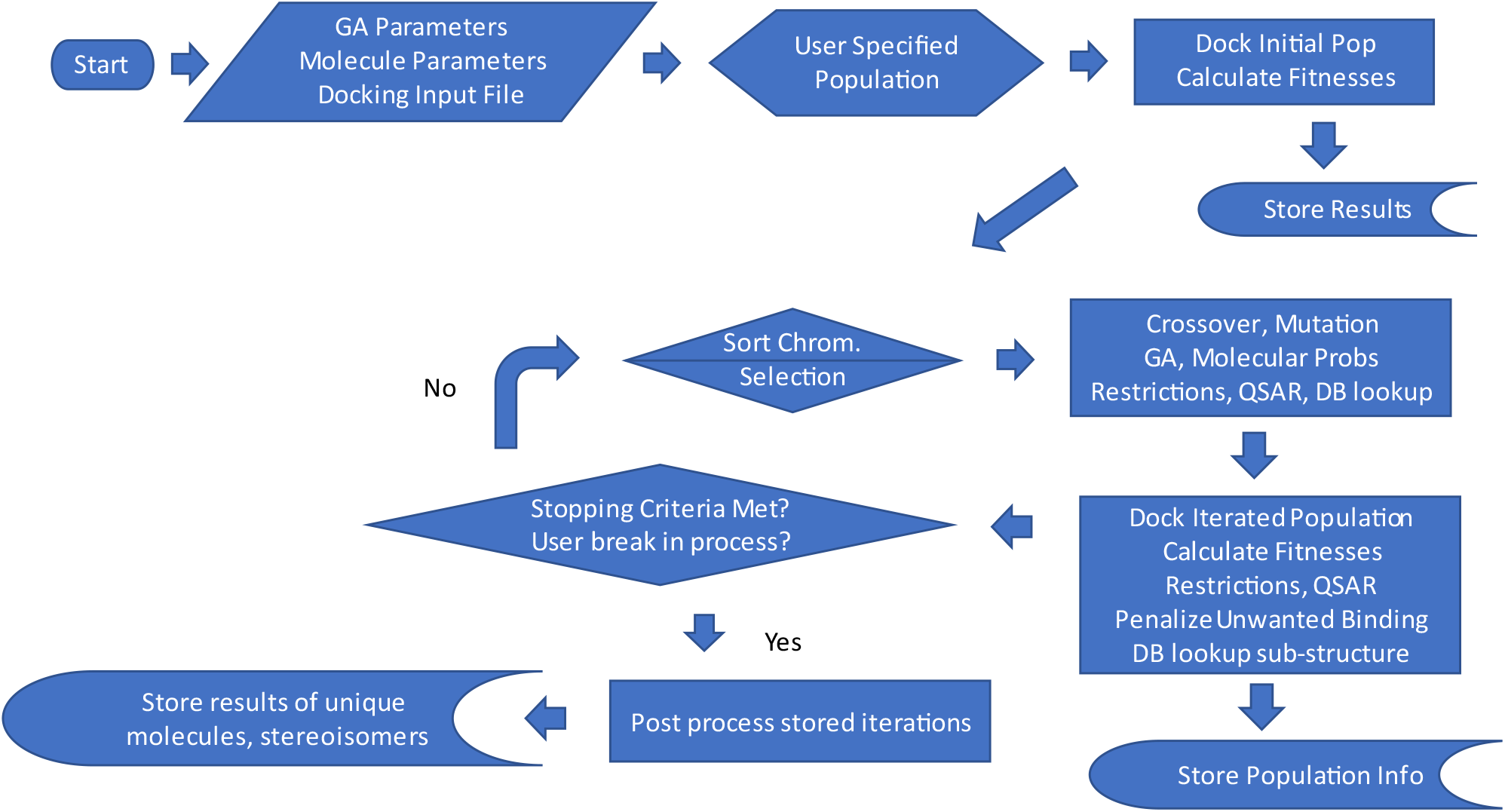
Flow of Ligand GA. With the default parameters, it is started with the inputs: ligand directory path, GOLD configuration file, output file name, initial population file name, and parameters. These are in the Ligand_GA_Config_File, prepared by the user (and defaulted).

The Ligand GA software is written in a very modular way. The functions are compact and documented. In all there are 40 files, most of them Matlab scripts, that are called in the execution of the head script. The software is designed to be very easy to use: the head function requires the input Global_Config file consisting of the ligand directory, the protein preparation conf file in CSD GOLD, the name of the output Matlab.dat file, and the initial GA molecule population file. If the program is terminated at any point, the output .dat file can be used to restart it from the termination point, as this .dat file has the information of the population at each iteration of the GA.

There are many necessary parameters used in the software. These have all been defaulted. However, any of them can and should be changed according to the molecular goal. These parameters and their context are listed in Tables 1 to 5. The point of listing these parameters in this article is to show the detail and flexibility of use in molecular construction. The various subfunctions and external path variables are also listed. The Ligand GA software package download has documentation of the various scripts and explanation of all of the parameters. Ligand GA does use the external software Corina Classic and CSD GOLD. The documentation of these are available online.

**Table 1:** GA parameters. These are the parameters for the operation of a Matlab GA in the global optimization toolbox with special crossover and mutation functions. These are defaults and the population size variable is calculated from the InitialPopFile input initial population file which consists of a column of SMILES strings. 12 parameters.

|  |  |  |  |  |
| --- | --- | --- | --- | --- |
| PopulationSize | pop_size |  | TolCon | .01 |
| InitialPopulation | Initial_pop |  | CrossoverFunction |  |
| Generations | 100 |  | MutationFunction |  |
| TolFun | .02 |  | OutputFcns |  |
| EliteCount | .1*pop_size |  | PlotFcn |  |
| StallGenLimit | 100 |  | Display | Iter |

There are two types of parameters required for the use: GA parameters and molecular parameters. Both sets have been defaulted for general usage but could be changed in specific protein and small molecule construction. These 54 parameters are listed in Tables I and II. The third set of 17 molecular ADME restriction parameters is given in Table 3. CSD GOLD requires a configuration file that has information about the prepared protein, the site, the GOLD GA parameters, and molecular chemical information relevant to the GOLD GA. Discussion of the use of CSD GOLD or Corina Classic is referred to in their extensive documentation.

The Ligand GA uses 13 mutation functions and 1 crossover function in an atypical fashion. Although the GA calls for a mutation or crossover after selection in the iteration process, it also randomly chooses one of the mutations out of the 13 in accordance with the molecular parameters. In addition, when a new atom is to be inserted into the chromosomal molecule or when an atom is to be changed into a different atom there are probabilities for choosing which atom to add or change to. These are also probabilities in the molecular parameter set following abundances of atoms in the molecular construction. These have been defaulted also in a uniform distribution. Note also that solely due to valence of the atoms, the set of which is to be chosen in the molecular parameter set (defaulted to C,N,O,S,P,F), there is a lean to more carbons (higher valence of 4) which has a chance of replacing a less valent atom then vice versa. Pseudo-atoms such as sulfates, phosphates, or any other segment can be included by specifying the symbol with valence, such as X=SO3 or PO3; multiple chars labeling have not been included yet in the different functions, only fundamental atom units, and will be in the form of 5 letter chains, such as X0001 or X0002.

### Tuning

A primary disadvantage of GA’s is the possible result in a local minimum of the objective function. This is common in any GA and particularly so with complicated fitness functions. Two ways, and there are many techniques, to avoid this are initialization of the population and scanning over crossover and mutation rates. Both have been included in the program. Initialization of the population could also be done by using the output of a previous run and selecting those individuals for diversity and non-trapping into a local minimum; this is a typical route for avoiding a local minimum.

Typically high GA crossover rates and low GA mutation rates of .8 and .2 are used in GA’s. Due to the number of different mutations and lack of sampling all of these mutations correctly in a limited number of iterations and no user input into the selection as the iterations continue, this GA application requires both GA rates high, .8 and .8 for example. Making and using a GA effectively requires 2 major parts: formulating the GA itself and tuning the GA parameters. Ligand GA, however, is not a typical GA due to the large number of molecule parameters listed in Table 2 and limited computing resources to find the best parameters. These molecule parameters, especially in the molecular mutation probabilities, also should be tuned to the specific target protein site and type of output small molecules. As a simple example, if the presence of rings is less wanted then the CLOSE_RING parameter in unnorm_mut_probability should be less than others, or if less branches are wanted then the ADD_BRANCH molecular mutation parameter should be less.

**Table 2:**
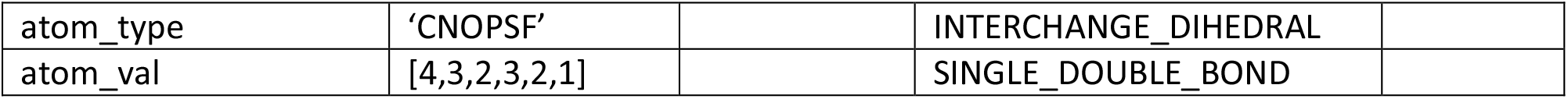
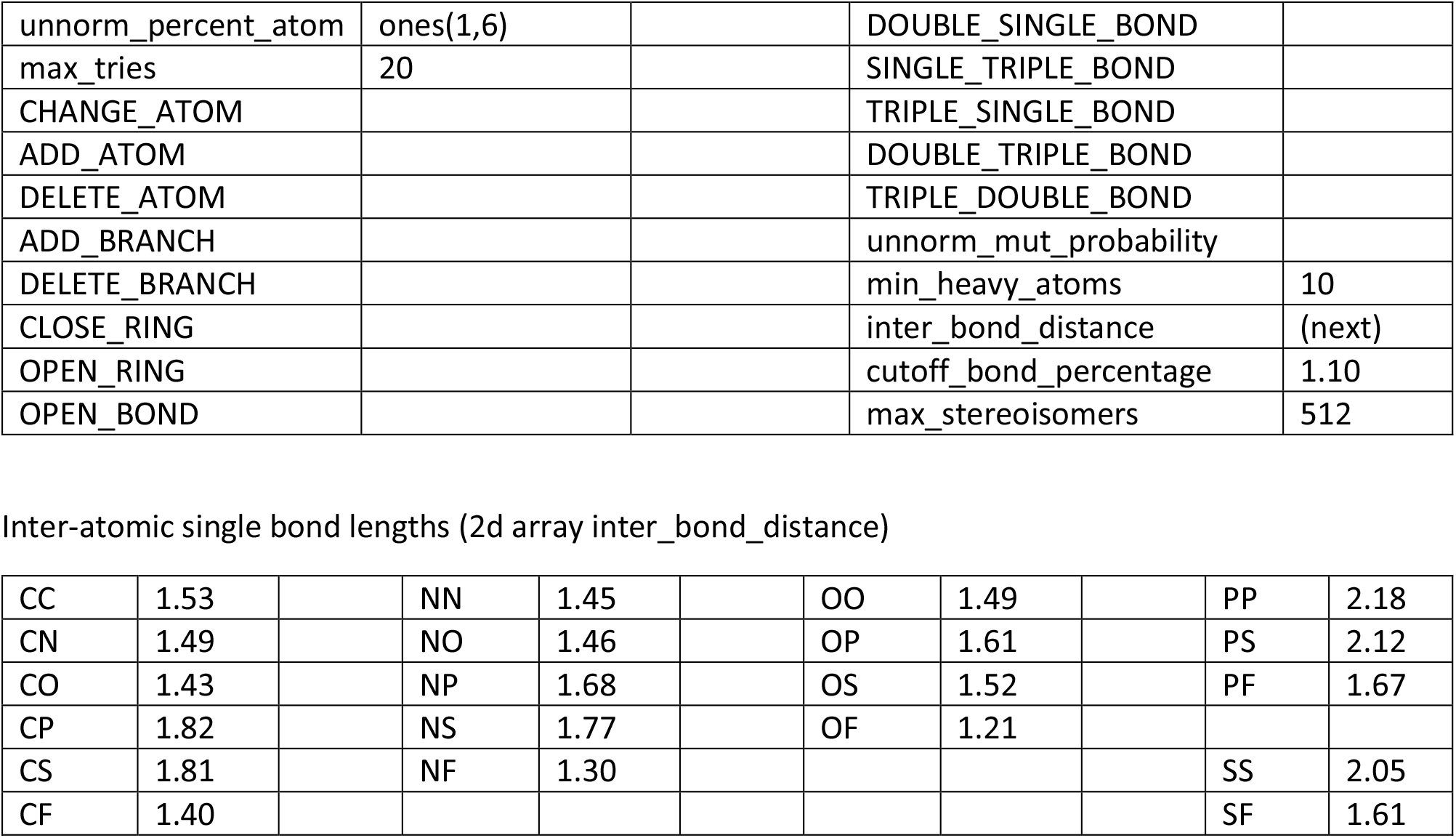
Molecular parameters. These molecular parameters are used to guide the chemical side of the Ligand GA evolution and are not internal GA parameters. These parameters are used to guide the evolution of the molecules and also specify the atomic content. 42 parameters.

|  |  |  |  |
| --- | --- | --- | --- |
| atom_type | 'CNOPSF' |  | INTERCHANGE_DIHEDRAL |
| atom_val | [4,3,2,3,2,1] |  | SINGLE_DOUBLE_BOND |

|  |  |  |  |  |
| --- | --- | --- | --- | --- |
| unnorm_percent_atom | ones(1,6) |  | DOUBLE_SINGLE_BOND |  |
| max_tries | 20 |  | SINGLE_TRIPLE_BOND |  |
| CHANGE_ATOM |  |  | TRIPLE_SINGLE_BOND |  |
| ADD_ATOM |  |  | DOUBLE_TRIPLE_BOND |  |
| DELETE_ATOM |  |  | TRIPLE_DOUBLE_BOND |  |
| ADD_BRANCH |  |  | unnorm_mut_probability |  |
| DELETE_BRANCH |  |  | min_heavy_atoms | 10 |
| CLOSE_RING |  |  | inter_bond_distance | (next) |
| OPEN_RING |  |  | cutoff_bond_percentage | 1.10 |
| OPEN_BOND |  |  | max_stereoisomers | 512 |

Inter-atomic single bond lengths (2d array inter\_bond\_distance)
|  |  |  |  |  |  |  |  |  |  |  |
| --- | --- | --- | --- | --- | --- | --- | --- | --- | --- | --- |
| CC | 1.53 |  | NN | 1.45 |  | OO | 1.49 |  | PP | 2.18 |
| CN | 1.49 |  | NO | 1.46 |  | OP | 1.61 |  | PS | 2.12 |
| CO | 1.43 |  | NP | 1.68 |  | OS | 1.52 |  | PF | 1.67 |
| CP | 1.82 |  | NS | 1.77 |  | OF | 1.21 |  |  |  |
| CS | 1.81 |  | NF | 1.30 |  |  |  |  | SS | 2.05 |
| CF | 1.40 |  |  |  |  |  |  |  | SF | 1.61 |

**Table 3:**
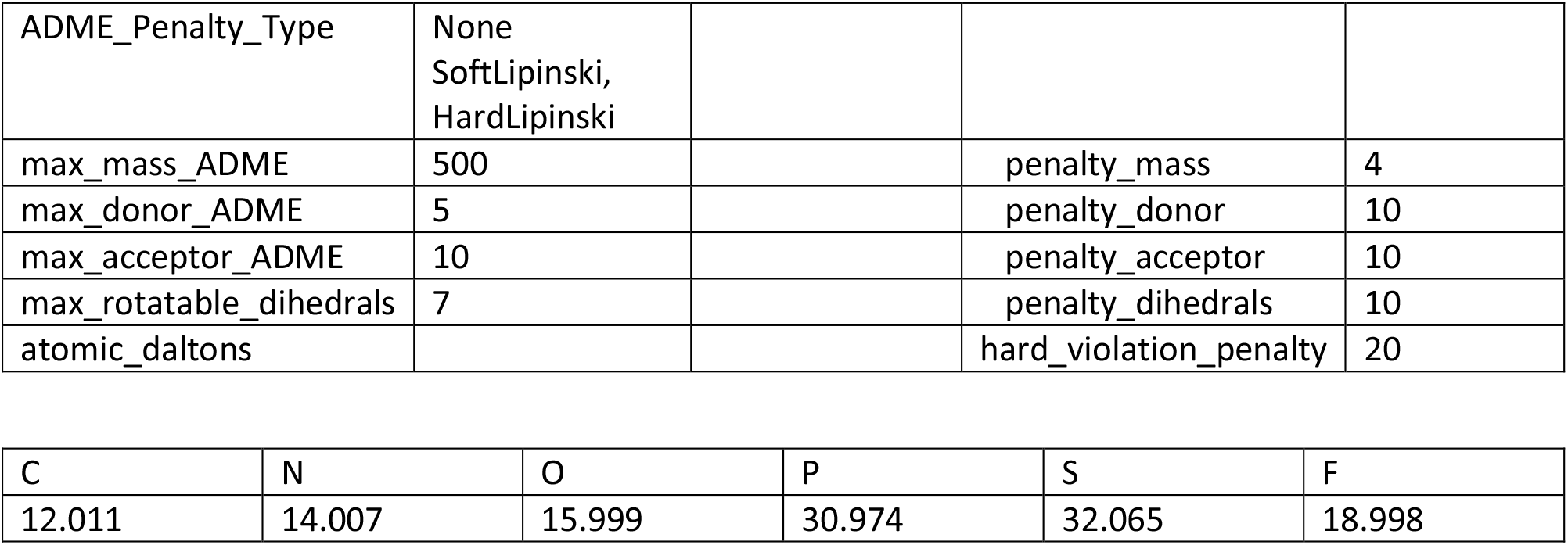
ADME restriction parameters. These are factors used in Lipinski’s Rule of 5 and subse uent rule of 3 (RO3). 17 parameters. Mass is in amu.

|  |  |  |  |  |
| --- | --- | --- | --- | --- |
| ADME_Penalty_Type | None<br>SoftLipinski,<br>HardLipinski |  |  |  |
| max_mass_ADME | 500 |  | penalty_mass | 4 |
| max_donor_ADME | 5 |  | penalty_donor | 10 |
| max_acceptor_ADME | 10 |  | penalty_acceptor | 10 |
| max_rotatable_dihedrals | 7 |  | penalty_dihedrals | 10 |
| atomic_daltons |  |  | hard_violation_penalty | 20 |

|  |  |  |  |  |  |
| --- | --- | --- | --- | --- | --- |
| C | N | O | P | S | F |
| 12.011 | 14.007 | 15.999 | 30.974 | 32.065 | 18.998 |

**Table 4:**
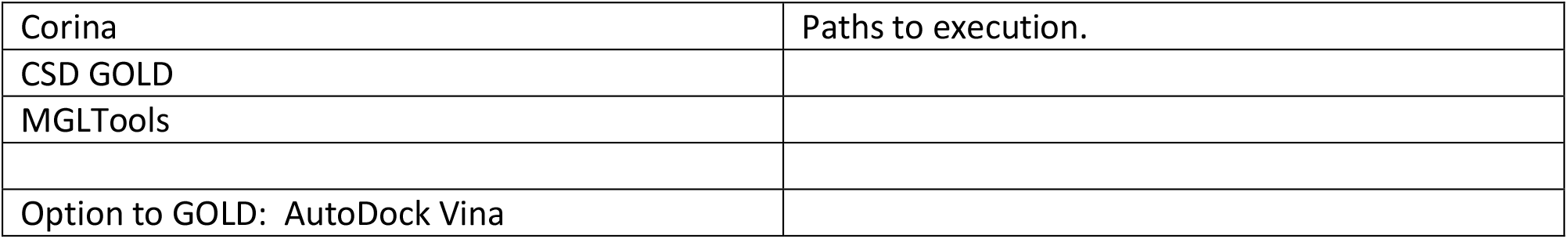
File paths to external software There are external programs used in Ligand GA. These path variables that need to be set in Ligand GA.

|  |  |
| --- | --- |
| Corina | Paths to execution. |
| CSD GOLD |  |
| MGLTools |  |
| Option to GOLD: AutoDock Vina |  |

**Table 5:** Software components. The head program Ligand GA and the various components used in its execution. 13 files.

|  |  |  |
| --- | --- | --- |
| Ligand_GA | Ligand_GA_Global | Ligand_GA_Output_Function |
| Ligand_GA_Crossover | Ligand_GA_Mutation | Ligand_GA_Load_Population |
| MolecularStructure | CLASH_CHECK | AtomCharVal |
| Ligand_GA_Fitness_Function_GOLD | RING_RENUMBER_CHECK | Ligand_GA_Pop_Save |
| Ligand_GA_Fitness_Function_Vina |  | Load_Population |

The last point requires more chemical information input in the choice of molecular mutation function; the mutation probabilities in a mutation is unnorm_mut_probability, which is defaulted to be uniform. These mutations guide the creation of atom change, addition or deletion of branches, opening or closing rings, and so on, and could be chosen differently depending on what type of structures in the output are wanted. This is difficult to guess before running the software, and in changing the mutation probabilities due to the complication of ligand protein interaction and docking. Sub-structure information in comparison with known molecules from a large database can alter the GA evolution towards more chemically desired molecules if this is included in the probabilistic choice of molecular mutations or other parameters.

### External software

Several external software packages are used in the operation of Ligand GA: Corina Classic, CSD GOLD, and MGLTools. Corina Classic is used to convert a non-isomeric SMILES string into a set of mol2 files describing its stereoisomers. GOLD is used for docking the small molecule to a potential binding site. MGLTools is used to convert a pdb file into an extended pdb file, that of a pdbqt file. This latter file has the explicit rotatable dihedral information; the pdbqt file is not strictly necessary as a short script can be used in its place to determine the rotatable bonds and branch information.

### Fitness function

The individual fitness is found by calling Corina to convert the non-isomeric SMILES chromosome to a set of stereoisomeric mol2 files and then calling the CSD GOLD docking program with the gold conf file, the latter is an input. There isn’t much freedom in the choices of fitness but there are different GOLD scores and PLP is used. A different score in this fitness can be used in changing the GOLD conf file.

This GOLD conf file has to be prepared before the start of Ligand GA, and it has a number of parameters and output choices to be used in the docking calculation. Two parameters important for computational efficiency are the popsiz and maxops. The example runs of modified Celecoxib and Simeprevir are computationally much harder than modified Aspirin due to the complexity increase of docking a designed drug and also in the number of stereoisomers; this can be alleviated by reducing the maxops in the GOLD GA to 10000 and in post-processing of particular molecule output using the default of 100000 to find more accurate docking poses. The GOLD output file gold_docking_results.txt has the results of a molecule’s docking to the binding site for the poses, and the best pose score is used after the fitness function script from parsing it. The parsing also extracts information about the hydrogen bonding. Further information from this file can be extracted and used in the molecular construction criteria; this would require adding more code to the fitness function.

The function Ligand GA Fitness Function GOLD takes an input of a cell array of non-isomeric SMILES strings. It firsts expands all of these molecules into their stereoisomers using Corina. The default limit on the max number is 512 (9 chiral centers), but can be changed with the parameter max_stereoisomers in the Ligand GA molecular parameter file. In the Ligand Dir there will be a directory created for each of the molecules given to the fitness function. Then, in each of the molecule sub-directories in ligand_dir there will be a sub-directory created for each of the molecule’s stereoisomers up to the maximum number. All of these directories are deleted in the call to Ligand GA Fitness Function GOLD and then created again for the molecules and stereoisomers. The output files from using GOLD on each stereoisomer docking is generated in the different stereoisomer sub-directories. File management is automated and there are automated safety precautions that except and not terminate the program if Corina couldn’t generate any mol2 files, if GOLD couldn’t find any docking poses, or if MGLTools couldn’t convert a pdb file to a pdbqt file; these instances are rare, but in principle could happen. If any of these happen a fitness of 0 is given to the non-isomeric SMILES chromosome and eventually it will be ejected from the population in the evolution.

### Constraints

The use of a GA in molecular construction has much flexibility. Penalties or constraints can be implemented in a variety of ways to guide the molecular evolution in the GA. Reliable ADME re uirements based on known heuristics such as Lipinski’s Rule of 5, Veber’s rule, the Ghose filter, and the Rule of 3 are necessary in drug candidate selection in the process of possible drug identification. These accurate heuristics for orally digested drugs (and in cases, for non-orally digested) can be implemented, with the exception of the octanol-water partition coefficient K_ow_ restraint, directly in the GA fitness function used in the molecular evolution. These requirements are from molecular weight, number of hydrogen bond donors and acceptors, number of rotatable dihedral bonds, and also polar surface area. All of these quantities for molecule and stereoisomer are calculated in the fitness evaluation if the parameter ADME_Penalty_Type is set to SoftLipinski or HardLipinski. These heuristics are then used in a soft or hard manner; hard means that the restriction is set by a cutoff with strict penalty and soft meaning that the restriction with a penalty set by the amount of violation. For example, requiring a maximal molecular mass of 500 could be set by (M_w_-500)*weight_penalty, by a strict M_w_<500, or by an overall large penalty if M_w_≥500.

In addition to ADME restrictions, binding aspects of the molecules can also be included in the GA evolution. The physical landscape of a protein or substrate surface has valleys, ridges, holes and is hilly. There may be equally optimal small molecule binders to the location of a target site, but when examined cover this surface in different ways although in the same vicinity. The requirement of the covering of this site in detail such as filling the valley or a hole can be included in the GA fitness function by including a penalty of physical localization of small molecule coordinates. A requirement of a valley or hole to be filled, or a ridge to be covered, can be imposed on the molecular evolution. This is important for the inhibition of a substrate in blocking the mechanism of interaction, for example.

The wanted presence of hydrogen bonds with particular amino acids can also be included as a restriction on the generated small molecules. Hydrogen bonding and their locations in the bound small ligand-protein complex is found in the GOLD output files, which are parsed to find the number of bonds and their amino acid or residue, and atomic location. A penalty can be included if a particular amino acid scale region of the surface is not covered or if a particular amino acid(s) does not share a hydrogen bond with the small molecule.

These restrictions are dependent on the protein. Only the ADME restrictions are included in this version of Ligand GA.

### Databases

As in conventional CADD, the inclusion of databases both in the evolution and also in the post-processing can also be included. Databases are commonly used for finding structure activity relationships and use a variety of QSAR methods. A database can be as simple as a list of SMILES strings with attached indexing to molecular information, such as the downloadable list from PubChem. The SMILES molecular format has been used extensively in estimating chemical properties of compounds based on sub-structure comparison with known molecules. Database searching can also be used for demanding a variety of physical constraints from the properties of known molecules.

An example of database usage is in the implementation of the K_ow_ ADME restriction, that log(P)<5 in Lipinski’s Rule of 5. This partition coefficient is difficult to compute, but can be guessed or estimated by comparing the molecule in the evolution with known molecules in a database that has log(P). During the fitness evaluation, this database can be called for substructure information to estimate the partition coefficient of the Ligand GA molecule during its evolution. Other purposes of including a database of known molecules include the use of potentially desired chemical properties of an evolving GA population of molecules.

The output of the Ligand GA generates a list of molecules in SMILES format and fitnesses. In the post-processing of this list, the structural similarities of these molecules can be compared with those in a list from an informed database. The textual SMILES format of the molecule is commonly used in pattern recognition in this manner. For example, a basic question such as, is this generated molecule structurally similar to any in a subset of the database?, can be answered. This information can be used in chemical synthesis or in potential FDA drug comparisons.

### Crossover and mutation

After the selection step, the crossovers and mutations are done internally in the GA. As mentioned there are a number of parameters involved. These are nonstandard functions and are listed in Table II. The molecular crossover will take the right side of a segment and interchange it with the right side of a different molecular segment. Segments are of 2 types An atom in a SMILES expression can be part of a branch, a closed ring, or neither. If the random atom selected in the crossover function is in a ring then the crossover will attempt again to find a segment; a ring cannot be split into 2 disjoint pieces by cutting one bond. If the atom is in a branch then the segment is from that atom to the end of the branch. If neither then the segment is from the atom to the end of the SMILES expression. The segments from the two selected parents are then interchanged. In the Ligand GA Global file there is a parameter, max_tries, which is the number of times to find a random segment of the parent molecule in the GA. It is defaulted to 20.

The mutation operators are self-explanatory by name and are commented and documented in the download.

### Selection

After the fitnesses are evaluated, the head program uses a defaulted selection process to determine those that are to be crossed over or mutated. The default is stochastic uniform but there are many GA choices such as roulette, tournament, and specialized selection functions. The selection process in the Matlab GA is not explained here but is referred to in the global optimization toolbox documentation [16].

### Storage of results

The population information at each iteration is stored in a Matlab data file that can be loaded into a Matlab session. It contains a cell array with the population of the molecules and the best fitnesses of these molecules. It does not include the stereoisomer individuals because non-isomeric SMILES is used, but this information of the molecule stereoisomers can be obtained by using the Ligand GA Fitness Function GOLD on any non-isomeric SMILES expression, or any set of molecules within a cell array input to this function.

### Output and post-processing

The script Ligand GA Output Function will take in the output .dat file and produce a sorted and unique list of molecules from the stored population information. The list can be used for a number of reasons, including the viability as a drug and the inspection in a medicinal chemical view towards synthesis. The list is also useful for examination of various biochemical properties and in comparison with existing databases such as PubChem. An example is to find out if any known and/or FDA approved drugs are structurally similar.

## Section 4: Discussion

A new software package, Ligand GA, is introduced that enables the construction of small molecules that are potential therapeutic or anti-viral drug candidates. The construction requires the information of the protein binding site and an initial set of small molecules. Ligand GA outputs lists of high scoring molecules in docking and binding which are designed according to multiple user criteria. The criteria can be very specific such as amino acid specific localization, ADME restriction, required hydrogen bonding, toxicity, and inclusion of database information.

Ligand GA has concise code which uses the textual non-isomeric SMILES format for the chromosomes and their manipulations. It also uses several proprietary external software: Matlab, Corina Classic, and CSD GOLD. The use of the software does not generate absolute results due to the nature of the genetic algorithm and the extremely complicated fitness (binding interactions of molecules), but it is useful for finding high affinity ligand binders. Ligand GA can be stopped during a run, then after an examination of the total output of the stored population from the iterations, can be restarted with a selection of these molecules either unchanged or manually edited. Running, restarting, running, … can improve results both in binding score and in types of molecules desired particular to the protein site amino acids desired to be chemically inhibited, amongst others.

Various examples of modified small drugs have been generated using this software including anti-inflammatory and anti-viral inhibitors. These generated molecules were examined for synthesizability, not included in Ligand GA. The software was designed for including geometric and molecular chemical constraints, and these can be included without changing the software. It was also written to interface with existing databases of molecules and drugs for input in the GA’s evolution and in using SMILES chromosomes in the post analysis.

The Ligand GA software, its structure, and the design methodology is explained. The software package download contains the Matlab code, documentation, and directories with the example inputs and outputs. There are instructions to use the software with minimal effort. The software is designed to expand by including constraints in the fitness function which are not included in this version.

Results from modifying Aspirin and Simeprevir are presented, in both of which there are thousands of generated molecules. These molecules illustrate the use of the software and also the design methodology, particularly in using constraints on the molecules. These molecules have higher docking and binding scores but also illustrate the specificity of the molecular construction in ADME restriction and localization within their corresponding protein binding sites.

The 30 CPU cluster typically has an extremely high but manageable load especially as the molecules evolve depending on their complexity. Scalability is built into Ligand GA. More computing resources and computing time enables a more thorough search and more specific results. Tuning the various parameters is also discussed.

[Note since 1^st^ submission: Ligand GA has been generalized to Ligand Multi-Protein GA and coded to use a multi-protein system: high binding/interaction of a small molecule to a target protein, low binding/interaction to a set of others, or any combination. This generalization was important in the conception of the project and is described in detail in ppt documents in the documentation folder in the download. This to be released version uses a multi-objective GA in minimizing more than one fitness function. As a first example the Pfizer Paxlovid/Ritonavir orally ingested therapeutic for Covid is being modified. Constructing an inhibitor of the SARS-Cov-2 3CL Mpro with reduced metabolic activity at the

CYP 3A4 enzyme (most common of the 6 liver CYP heme iron carrying enzymes) would eliminate the need for Ritonavir after modifications from an initial starting point of Paxlovid. This example use is in progress with positive partial results. In going from 1 protein to an N-protein system, the computational requirements multiply by roughly a factor of N, but also depends on the complication of the molecule and protein binding sites. The use of the Georgia Advanced Computing Resource Center (GACRC) Sapelo2 cluster and their hundreds of nodes with modern cores is appreciated.]

## Acknowledgements

G.C. thanks Christian Heiss and James H. Prestegard for useful conversations, Robert J. Woods for the use of a 30 CPU cluster, and the Complex Carbohydrate Research Center for hospitality.

## Statements and declarations

There are no conflicts of interest.

## Software download

https://www.dropbox.com/sh/s5nm3yzsd3l4y5r/AADCjHIymuu0nWSYnD3V0dQRa?dl=0

The download contains the software, instructions and documentation, files used in the paper, and several example directories. The Previous Calcs directories have .mat, not .m, files which are large in memory and do not have to be downloaded. There are 2 .tar.gz files, one with the .mat files and one without, of 87 and 11 MB. The output directories of a full length GOLD calculation (not efficient, but good for accurate docking scores and used in the paper) of a non-isomeric molecule expression can be large, in excess of 1 GB and not included in the downloads.

Contact for the files if there is an issue. (also, https://github.com/GordonChalmers/GRC_Presentations)

## Footnotes

1 An initial examination of 4 heavy atom types with valence greater than 1 in a linear chain gives 4^*N*^, with *N* the number of heavy atoms. The exact number of molecules given atomic and pseudo-residue content can be calculated from graph counting developments in early and current studies of large orders in perturbation theory of quantum field theories.

2 This software is extensively documented. Stereoisomer and ring conformation information is also in the output.

3 There are many different SMILES expressions for a given molecule. The IUPAC convention for unique SMILES is not implemented in this work. Due to the multiple paths from molecule A to molecule B in modifications, a unique representation may restrict the search, although conversion to a unique form is useful in database searching.

4 The details of CSD GOLD and its use are documented at [15].

5 The choice of a soft limit that is a penalty proportional to the violation or a direct hard limit can have an effect on the GA evolution of the population.

## References

1. Marsland, S. Machine Learning, An Algorithm Perspective (2 ed.). Chapman and Hall, CRC. 2015, 430 pp. DOI:10.1201/9781420067194

2. Eberhardt RC, Shi RC. Computational Intelligence: Concepts to Implementations. Morgan Kauffman, Elsevier, 2007, 496 pp. DOI:10.1016/B978-155860759-0/50009-3. ISBN: 978-1-55860-759-0.

3. Goldberg DE. Genetic Algorithms in Search, Optimization, and Machine Learning (13 ed.). Addison-Wesley, 1988, 432 pp. ISBN-13: 978-0201157673

4. Dumitrescu D, Beatrice Lazzerini B, Jain LC, Dumitrescu A. Evolutionary Computation. CRC Press. 1^st^ Ed., 2000, 424 pp. DOI: 10.1201/9781482273960

5. European Consortium. (2021). Exscalate4Cov. Retrieved from EU Exscalate 4Cov Project: https://www.exscalate4cov.eu/index.html

6. PubChem. (2021). Retrieved from National Center for Biotechnology Information, National Institutes of Health: https://pubchem.ncbi.nlm.nil.gov

7. Sterling T, Irwin JJ. ZINC 15—Ligand Discovery for Everyone. Journal of Chemical Information and Modeling 2015, 55(11):2324–2337. DOI: 10.1021/acs.jcim.5b00559

8. Irwin JJ, Sterling T, Mysinger MM, Bolstad ES, Coleman RG. ZINC: A free tool to discover chemistry for biology. Chem. Inf. Model. 2012, 52(7), 1757–1768. DOI:10.1021/ci2001277

9. Sterling T, Irwin JJ. ZINC -- a free database of commercially available compounds for virtual screening. J. Chem. Inf. Model 2012, 45(1), 177–82. DOI:10.1021/ci049714

10. Zinc database. (2021). Retrieved from Zinc: http://zinc.docking.org/

11. Wishart DS, Feunang YD, Guo AC, Lo EJ, Marcu A, Grant JR, Sajed T, Johnson D, Li C, Sayeeda Z, Assempour N, Iynkkaran I, Liu Y, Maciejewski A, Gale N, Wilson A, Chin L, Cummings R, Le D, Pon A, Knox C, Wilson M. DrugBank 5.0: a major update to the DrugBank database for 2018. Nucleic Acids Res. 2017 Nov 8. PubMed: 29126136 doi: 10.1093/nar/gkx1037.

12. Law V, Knox C, Djoumbou Y, Jewison T, Guo AC, Liu Y, Maciejewski A, Arndt D, Wilson M, Neveu V, Tang A, Gabriel G, Ly C, Adamjee S, Dame ZT, Han B, Zhou Y, Wishart DS. DrugBank 4.0: shedding new light on drug metabolism. Nucleic Acids Res. 2014, 42(1):D1091-7. PubMed: 24203711

13. Knox C, Law V, Jewison T, Liu P, Ly S, Frolkis A, Pon A, Banco K, Mak C, Neveu V, Djoumbou Y, Eisner R, Guo AC, Wishart DS. DrugBank 3.0: a comprehensive resource for ‘omics’ research on drugs. Nucleic Acids Res. 2011, 39(Database issue):D1035–41. PubMed: 21059682

14. Wishart DS, Knox C, Guo AC, Cheng D, Shrivastava S, Tzur D, Gautam B, Hassanali M. DrugBank: a knowledgebase for drugs, drug actions and drug targets. Nucleic Acids Res. 2008, 36(Database issue):D901–6. PubMed: 18048412

15. Wishart DS, Knox C, Guo AC, Shrivastava S, Hassanali M, Stothard P, Chang Z, Woolsey J. DrugBank: a comprehensive resource for in silico drug discovery and exploration. Nucleic Acids Res. 2006, 34(Database issue):D668–72. PubMed: 16381955

16. DrugBank Online. (2021). Retrieved from DrugBank Online: https://go.drugbank.com/

17. Cambridge Structural Database (CSD), Cambridge Crystallographic Data Centre. (2021). Retrieved from CSD: http://www.ccdc.cam.uk/solutions/csd-core/components/csd/

18. Royal Society of Chemistry. (2021). ChemSpider. Retrieved from http://www.chemspider.com

19. ChEMBL, EMBL’s European Bioinformatics Institute (EMBL-EBI). (2021). Retrieved from ChEMBL: https://www.ebi.ac.uk/chembl/

20. Mendez D, Gaulton A, Bento AP, Chambers J, De Veij M, Félix E, Magariños MP, Mosquera JF, Mutowo P, Nowotka M. ChEMBL: towards direct deposition of bioassay data. Nucleic Acids Res. 2019, 47(D1), D930:D940. DOI: 10.1093/nar/gky1075 PMID: 25883136

21. Davies M, Nowotka M, Papadatos G, Dedman N, Gaulton A, Atkinson F, Bellis L, Overington JP. ChEMBL web services: streamlining access to drug discovery data and utilities Nucleic Acids Res. 2015, 43(Web Server issue): W612–W620. DOI: 10.1093/nar/gkv352 PMID: 25883136

22. Weininger, D. SMILES, a chemical language and information system. 1. Introduction to methodology and encoding rules. J. Chem. Inf. Comput. Sci. 1988, 28(1), 31–36. doi:10.1021/ci00057a005

23. Weininger D, Weininger A, Weininger J. SMILES. 2. Algorithm for generation of unique SMILES notation. J. Chem. Inf. Comput. Sci. 1998, 29(2), 97–101. doi:10.1021/ci00062/a008

24. Weininger D. Smiles. 3. Depict. Graphical depiction of chemical structures. J. Chem. Inf. Comput. Sci. 1990, 30(3), 237–243. doi:10.1021/ci00067a005

25. D.A. Case, H.M. Aktulga, K. Belfon, I.Y. Ben-Shalom, S.R. Brozell, D.S. Cerutti, T.E. Cheatham, III, G.A. Cisneros, V.W.D. Cruzeiro, T.A. Darden, R.E. Duke, G. Giambasu, M.K. Gilson, H. Gohlke, A.W. Goetz, R. Harris, S. Izadi, S.A. Izmailov, C. Jin, K. Kasavajhala, M.C. Kaymak, E. King, A. Kovalenko, T. Kurtzman, T.S. Lee, S. LeGrand, P. Li, C. Lin, J. Liu, T. Luchko, R. Luo, M. Machado, V. Man, M. Manathunga, K.M. Merz Miao, O. Mikhailovskii, G. Monard, H. guyen, K.A. O’Hearn, A. Onufriev, F. Pan, S. Pantano, R. iA. Rahnamoun, D.R. Roe, A. Roitberg, C. Sagui, S. Schott-Verdugo, J. Shen, C.L. Simmerling, N.R. Skrynnikov, J. Smith, J. Swails, R.C. Walker, J. Wang, H. Wei, R.M. Wolf, X. Wu, Y. Xue, D.M. York, S. Zhao, and P.A. Kollman (2021), Amber 2021, University of California, San Francisco.

26. Sadowski J, Gasteiger J., Klebe G. Comparison of Automatic Three-Dimensional Model Builders Using 639 X-Ray Structures. J. Chem. Inf. Comput. Sci. 1994, 34, 1000–1008 DOI: 10.1021/ci00020a039

27. Schwab CH. Conformations and 3D pharmacophore searching. Drug Discovery Today: Technologies 2010, 7(4), Winter 2010, e245–e253 DOI: 10.1016/j.ddtec.2010.10.003

28. Molecular Networks GmbH, Altamira, LLC. (2021). Corina. Retrieved from MN-AM Corina: https://www.mn-am.com/products/corina

29. Schwab CH. Molecular Structure Representation in Chemoinformatics Applications, Schwab CH, BigChem Autumn 2017 School, Modena, Italy. Received from http://bigchem.eu/sites/default/files/School3_Schwab.pdf

30. Jones G, Willett P, Glen RC, Leach AR, Taylor R. (1997). Development and validation of a genetic algorithm for flexible docking. J. Mol. Biol. 1997, 267(3), 727–748. DOI: 10.1006/jmbi.1996.0897 PMID: 9126849

31. Cambridge Crystallographic Data Centre. (2021). CCDC Discovery GOLD. Retrieved from GOLD Protein Ligand Docking Software: https://www.ccdc.cam.ac.uk/solutions/csd-discovery/Components/Gold/

32. MathWorks, Inc. MATLAB. (2020b). Retrieved from MathWorks: https://www.mathworks.com/

33. Molecular Graphics Laboratory at the Scripps Research Institute. (2021). MGL Tools. Retrieved from MGLTools: http://mgltools.scripps.edu/

34. Trott O, Olsen AJ. AutoDock Vina: improving the speed and accuracy of docking with a new scoring function, efficient optimization and multithreading. Journal of Computational Chemistry 2010, 31(2), 455–461. DOI: 10.1002/jcc.21334

35. Molecular Graphics Lab at The Scripps Research Institute. (2021). AutoDock Vina. Retrieved from AutoDock Vina: http://vina.scripps.edu/

36. Pagadala NS, Syed K, Tuszynski J. Software for molecular docking: a review. Biophys. Rev. 2017, 9(2):91–102. DOI: 10.1007/s12551-016-0247-1 PMID: 28510083 PMCID: PMC5425816

37. Plewc ynski D, Łaźniewski M, Augustyniak R, Ginalski K. Can we trust docking results Evaluation of seven commonly used programs on PDBbind database comparative study. J. Comput. Chem. 2011, 32(4):742–55. DOI: 10.1002/jcc.21643 PMID: 20812323

38. Durrant JD, Amaro R, McCammon JA, AutoGrow: A novel algorithm for protein inhibitor design, Chem. Bio. Drug Design 2009, 73(2):168–178. DOI: 10.1111/j.1747-0285.2008.00761.x

39. Spiegel JO, Durrant JD, AutoGrow4: an open source genetic algorithm for de novo drug design and lead optimization, J. Cheminform. 2020, 12:25. DOI: 10.1186/s13321-020-00429-4

40. Durrant JD, Lindert S, McCammon JA, AutoGrow 3.0: An improved algorithm for chemically tractable, semi-automatic protein inhibitor design, J. Mol. Graph and Modelling 2013, 44:104–112. DOI: 10.1016/j.jmgm.2013.05.006

41. Kawai K, Nagata N, Takahashi Y, De novo design of drug-like molecules by a fragment-based molecular evolutionary approach, J. Chem. Inf. Model. 2014, 54(1):49–56. DOI: 10.1021/ci400418c

42. Lipinski CA, Lombardo F, Dominy WB, Feeney PJ. Experimental and computational approaches to estimate solubility and permeability in drug discovery and development settings. Adv. Drug Deliv. Rev. 2001, 46(1-3), 3–26. DOI: 10.1016/s0169-409x(00)00129-0

43. Lipinski C. Lead-and drug-like compounds: the rule-of-five revolution. Drug Discovery Today Technologies 2004, 1(4), 337–341. PMID: 24981612 DOI: 10.1016/j.ddtec.2004.11.007

44. Ghose AK, Viswanadhan VN, Wendoloski JJ. A knowledge-based approach in designing combinatorial or medicinal chemistry libraries for drug discovery. J. Comb. Chem. 1999, 1(1), 55–68. DOI: 10.1021/cc9800071.

45. Veber DF, Johnson SR, Cheng HY, Smith BR, Ward KW, Kobble KD. Molecular properties that influence the oral bioavailability of drug candidates. J. Med. Chem. 2002, 45(12), 2615–23. PMID: 12036371 DOI: 10.1021/jm020017n

46. Congreve M, Carr R, Murray C, Jhoti H. A ‘rule of three’ for fragment-based lead discovery? Drug Discov. Today 2003, 8(19), 876–877. DOI: 10.1016/S1359-6446(03)02831-9

47. Benet LZ, Hosey CM, Ursu O, Oprea TI. BDDCS, the Rule of 5 and Drugability. Adv Drug Deliv Rev. 2016, 101, 89–98. DOI: 10.1016/j.addr.2016.05.007

48. Lucido MJ, Orlando BJ, Vecchio AJ, Malkowski MG. Crystal Structure of Aspirin-Acetylated Human Cyclooxygenase-2: Insight into the Formation of Products with Reversed Stereochemistry. Biochemistry 2016, 55, 1226–1238. DOI: 10.1021/acs.biochem.5b01378 PMID: 26859324

49. Orlando BJ, Lucido MJ, Malkowski MG. The structure of Ibuprofen bound to cyclooxygenase-2. J. Struct. Biol. 2015, 189, 62–66. DOI:10.1016/j.jsb.2014.11.005

50. Pettersen EF, Goddard GT, Huang CC, Couch GS, Greenblatt DM, Meng EC, Ferrin TE. UCSF Chimera--a visualization system for exploratory research and analysis. J. Comput. Chem. 2004, 25(13), 1605-PMID: 15264254 DOI: 10.1002/jcc.20084

51. University of California at San Francisco (UCSF) -Resource for Biocomputing, V. a. (Current). UCSF Chimera, an Extensible Molecular Modeling System. Retrieved from UCSF Chimera: https://www.cgl.ucsf.edu/chimera/

52. Humphrey W, Dalke A, Schulten K. VMD - Visual Molecular Dynamics. J. Molec. Graphics 1996, 14(1), 33–38. DOI: 10.1016/0263-7855(96)00018-5

53. University of Illinois at Urbana-Champaign, Theoretical and Computational Biophysics Group. (2021). VMD - Visual Molecular Dynamics. Retrieved from VMD - Visual Moleular Dynamics: https://www.ks.uiuc.edu/Research/vmd/

54. The PyMOL Molecular Graphics System, Version 2.0 Schrödinger, LLC. Retrieved from PyMOL: http://www.pymol.org/pymol

55. Kingsley LJ, Brunet V, Lelais G, McCloskey S, Milliken K, Leija E, Fuhs SR, Wang K, Zhou E, Spraggon G. (2019). Development of a virtual reality platform for effective communication of structural data in drug discovery. Journal of Molecular Graphics and Modeling 2019, 89, 234–242. DOI:10.1016/j.jmgm.2019.03.010

56. Nanome, Inc. (2021). Nanome. Retrieved from Nanome: https://nanome.ai/

57. Simeprevir. Retrieved from Pubchem: https://pubchem.ncbi.nlm.nih.gov/compound/simeprevir

58. Anson B, Mesecar A. (2020). X-ray structure of SARS-Cov-2 main protease bound to boceprevir at 1.45 A. Published online. doi:10.2210/pdb6WNP/pdb

59. Bafna K, Krug RM, Montelione GT. (2020). Structural Similarity of SARS-CoV2 M pro and HCV NS3/4A Proteases Suggests New Approaches for Identifying Existing Drugs Useful as COVID-19 Therapeutics. DOI: 10.26434/chemrxiv.12153615. Preprint. PMID: 32511291 PMCID: PMC7263768

60. Bafna K, White K, Harish B, Rosales R, Ramelot TA, Acton TB, Moreno E, Kehrer T, Miorin L, Royer CA, García-Sastre A, Krug RM, Montelione GT. Hepatitis C virus drugs that inhibit SARS-CoV-2 papain-like protease synergize with remdesivir to suppress viral replication in cell culture. Cell Rep. 2021, 35(7):109133. DOI: 10.1016/j.celrep.2021.109133, PMID: 33984267, PMCID: PMC8075848

61. Matthew D Hall, James M Anderson, Annaliesa Anderson, David Baker, Jay Bradner, Kyle R Brimacombe, Elizabeth A Campbell, Kizzmekia S Corbett, Kara Carter, Sara Cherry, Lillian Chiang, Tomas Cihlar, Emmie de Wit, Mark Denison, Matthew Disney, Courtney V Fletcher, Stephanie L Ford-Scheimer, Matthias Götte, Abigail C Grossman, Frederick G Hayden, Daria J Hazuda, Charlotte A Lanteri, Hilary Marston, Andrew D Mesecar, Stephanie Moore, Jennifer O wankwo, Jules O’Rear, George Painter, Kumar Singh Saikatendu, Celia A Schiffer, Timothy P Sheahan, Pei-Yong Shi, Hugh D Smyth, Michael J Sofia, Marla Weetall, Sandra K Weller, Richard Whitley, Anthony S Fauci, Christopher P Austin, Francis S Collins, Anthony J Conley, Mindy I Davis. The Journal of Infectious Diseases 2021, 224(S1)1:S1–S21. https://doi.org/10.1093/infdis/jiab305

62. PDBe-KB COVID-19 Data Portal at Protein Data Bank in Europe. (2021). Retrieved from: https://www.ebi.ac.uk/pdbe/covid-19

